# The Grainyhead/LSF transcription factor GRH-1 is rhythmically required for molting

**DOI:** 10.1101/2022.03.01.482504

**Authors:** Milou W.M. Meeuse, Yannick P. Hauser, Smita Nahar, Kathrin Braun, Helge Großhans

## Abstract

Molting, that is, the synthesis and shedding of a cuticular exoskeleton, is a defining characteristic of ecdysozoa. In nematodes such as *C. elegans*, molts rhythmically terminate each of four larval stages. The molting cycle is tightly coupled to the rhythmic accumulation of thousands of transcripts. Here, using chromatin immunoprecipitation coupled to sequencing (ChIP-seq) and quantitative reporter assays, we show that these dynamic gene expression patterns rely on rhythmic transcription. To gain insight into the relevant gene regulatory networks (GRNs), we performed an RNAi-based screen for transcription factors required for molting to identify potential components of a molting clock. We find that depletion of GRH-1, BLMP-1, NHR-23, NHR-25, MYRF-1 or BED-3 impairs progression through the molting cycle. We characterize GRH-1, a Grainyhead/LSF transcription factor whose orthologues in other animals are key epithelial cell fate regulators. We show that GRH-1 depletion causes a dose-dependent extension of molt duration, defects in cuticle formation and shedding, and larval death. Coincident with its rhythmic accumulation, GRH-1 is required repetitively for each molt, during specific time windows preceding lethargus. These findings are consistent with a function of GRH-1 in a molting cycle GRN. As its mammalian orthologues, as well as those of BLMP-1 and NHR-23, have been implicated in rhythmic homeostatic skin regeneration in mouse, the mechanisms underlying rhythmic *C. elegans* molting may apply beyond nematodes.

## Introduction

*C. elegans* larval development subdivides into four larval stages, each terminated by a molt. In a first step, termed apolysis, the connections of the existing cuticle to the underlying epidermis are severed. Subsequently, a new cuticle is synthesized, before the old cuticle is shed in a final step termed ecdysis. Traditionally, the time of molting is equated with lethargus, a period of relative behavioral quiescence, when animals stop feeding. For simplicity we will follow this tradition here, but note that additional events required for successful molting precede lethargus (Cohen et al., 2020; Cohen and Sundaram, 2020; Tsiairis and Großhans, 2021).

Molts occur at regular intervals of 7 – 8 hr at 25°C, and a clock-type mechanism has been invoked to explain this regularity (Monsalve and Frand, 2012; Tsiairis and Großhans, 2021). Such a clock mechanism may also explain, and be partially based on, the rhythmic expression of thousands of genes that is coupled to the molting cycle (Hendriks et al., 2014; Kim et al., 2013; Meeuse et al., 2020; Turek and Bringmann, 2014). However, the components of this clock, and accordingly their wiring, have remained largely elusive (Tsiairis and Großhans, 2021).

Counter-intuitively, rhythmic mRNA accumulation in the mammalian circadian clock appears to rely chiefly on co-and post-transcriptional mechanisms, including rhythmic splicing (Koike et al., 2012; Menet et al., 2012; Preußner et al., 2017). Here, we demonstrate that transcript level oscillations in *C. elegans* larvae are parsimoniously explained by rhythmic RNA polymerase II recruitment to promoters. This finding suggests that rhythmically active transcription factors are components of the underlying machinery, or core oscillator, and thus presumably also the molting cycle clock. To identify possible candidates, we screened through a selection of rhythmically expressed transcription factors, assuming that rhythmic transcription would be a possible (though not necessarily the only) mechanism of achieving rhythmic transcription factor activity. From a set of 92 such transcription factors (Hendriks et al., 2014), we identified six whose depletion altered molt number and/or duration. These include the nuclear hormone receptors NHR-23 and NHR-25 and the myelin regulatory family transcription factor MYRF-1/PQN-47, which had previously been linked to molting (Frand et al., 2005; Gissendanner et al., 2004; Gissendanner and Sluder, 2000; Johnson et al., 2021; Kostrouchova et al., 1998; Kostrouchova et al., 2001; Meng et al., 2017; Russel et al., 2011), as well as novel factors.

We characterize the function of GRH-1, the sole *C. elegans* member of the phylogenetically conserved LSF/Grainyhead family (Venkatesan et al., 2003). Grainyhead proteins are key regulators of differentiation, maintenance, integrity and repair of different epithelial tissues in animals (Sundararajan et al., 2020). RNAi-mediated depletion of *C. elegans* GRH-1 causes embryonic death, potentially due to cuticular defects (Venkatesan et al., 2003). We report that post-embryonically, GRH-1 accumulates rhythmically and promotes molting through its activity in a specific window during each larval stage. Its depletion delays the onset of ecdysis in a dose-dependent manner to the point that animals severely depleted arrest development and die by bursting through a defective cuticle.

Our results, together with the validation of an additional hit, BLMP-1, in separate work by us and others (Hauser et al., 2021; Stec et al., 2021), provide new insights into the transcriptional mechanisms that support rhythmic molting and identify potential molting clock components. The fact that GRH-1 as well as additional screen hits also function in rhythmic homeostatic skin regeneration in mouse (Magnusdottir et al., 2007; Steinmayr et al., 1998; Telerman et al., 2017; Wilanowski et al., 2008), suggests mechanistic similarities between this process and the molting process of nematodes.

## Results

### Rhythmic transcription of oscillating genes is driven by rhythmic RNA Polymerase II occupancy

Previous observations that the levels of intronic RNA encoded by oscillating genes also oscillate (Hendriks et al., 2014) provided circumstantial evidence for a model of rhythmic gene transcription. However, technical limitations restricted this analysis to a set of highly expressed genes with long introns, and genuine pre-mRNAs could not be distinguished from excised introns. Hence, we employed temporally resolved DNA-dependent RNA polymerase II (RNAPII) chromatin immunoprecipitation coupled to sequencing (ChIP-seq) to examine the dynamics of RNAPII binding to oscillating gene promoters. We used synchronized wild-type worms collected hourly from 22 hours until 33 hours of post-embryonic development at 25°C and quantified a 1-kb window around the transcription start sites (TSSs) as a proxy for temporal RNAPII promoter occupancy on annotated oscillating genes (Meeuse et al., 2020). This revealed widespread rhythmic binding of RNAPII at the promoters of genes for which we also observed mRNA level oscillations in RNA sequencing data generated from the same samples (**Figure 1A,B, Figure S1**).

**Fig 1:**
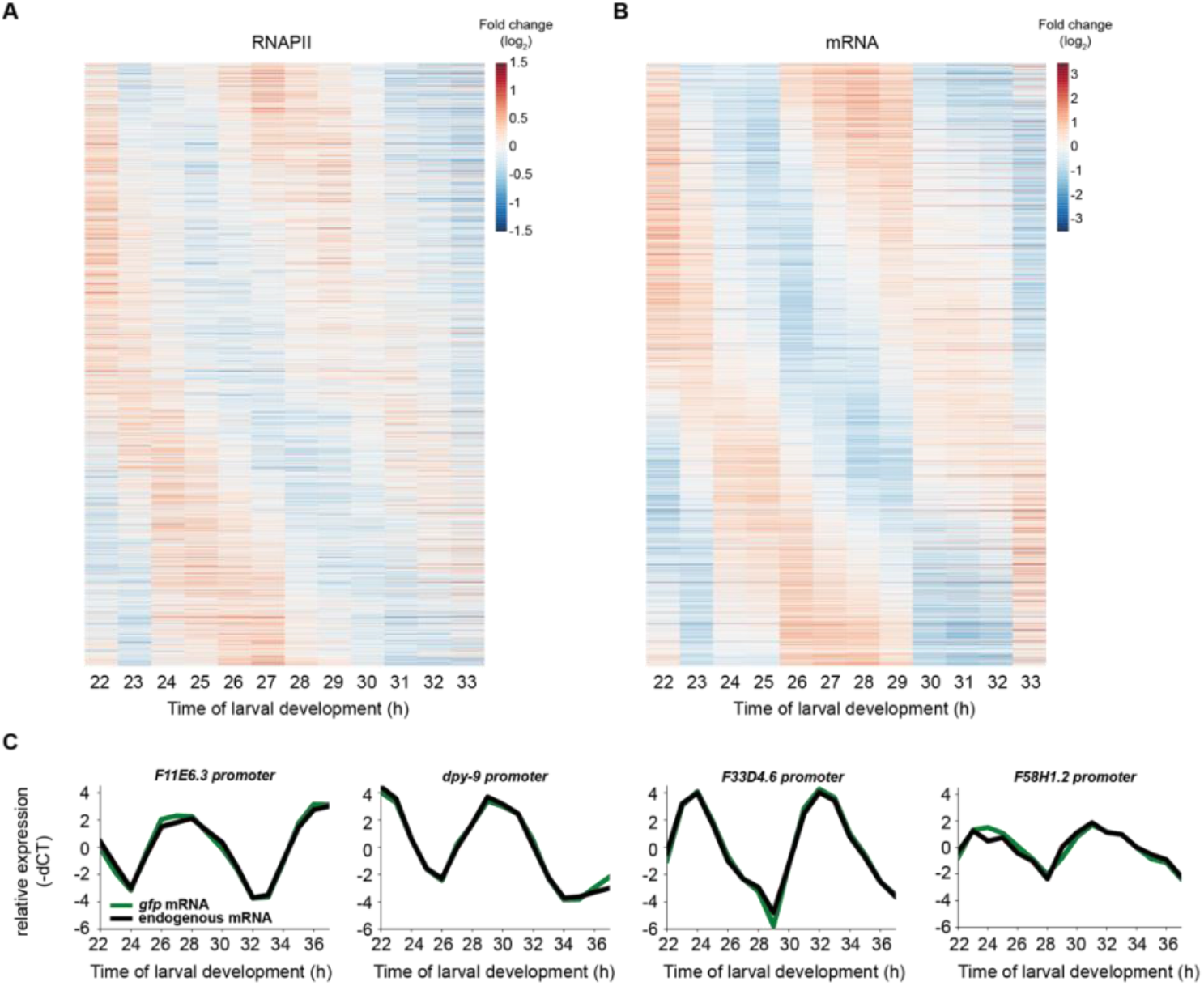
Oscillatory gene expression arises from promoter-driven rhythmic transcription. **A, B**, Log_2_-transformed, mean-normalized RNA polymerase II ChIP (A) and RNA (B) sequencing reads of oscillating genes detectably expressed at the chosen sequencing depth (n=2,106). Reads are ordered according to peak phase obtained from Meeuse et al., 2020. **C**, RT-qPCR time courses. Promoters of the indicated oscillating genes were used to drive expression of a destabilized, nuclear GFP from a single copy integrated transgene; *gfp* mRNA and the endogenous transcript driven by the same promoter were quantified from the same RNA samples. Relative expression was calculated as (target Ct values – actin Ct values) * (−1) and then mean normalized for each trace individually. See also **Figures S1, S2**.

We also detected instances where oscillating mRNA levels were not accompanied by rhythmic RNAPII promoter binding. It is possible that this might reflect instances of post-transcriptional regulation. However, we consider this more likely a technical limitation, because we noticed a general reduction of amplitudes in the ChIP-sequencing relative to the RNA-sequencing experiment, probably reflecting a reduced dynamic range of the former over the latter (see also below). This limitation notwithstanding, the data support the notion that rhythmic transcription is a major contributor to rhythmic transcript accumulation and specifically point to rhythmic recruitment of RNAPII as a relevant mechanism.

### Promoter-driven *gfp* reporter transgenes recapitulate transcription of endogenous genes

To understand in more detail how rhythmic transcription drives mRNA level oscillations, we characterized transcriptional reporters that contained the putative promoters (either 2kb upstream of the ATG or until the next upstream gene) of oscillating genes fused to a sequence encoding destabilized, nucleus-localized green fluorescent protein (GFP). We chose promoters from genes with a variety of peak expression phases. To exclude 3’UTR-mediated posttranscriptional regulation, the reporters further contained the *unc-54* 3’UTR, which we selected because *unc-54* did not display transcript level oscillation in our mRNA sequencing time courses (Hendriks et al., 2014; Meeuse et al., 2020) and because its 3’UTR appears devoid of regulatory activity (Brancati and Grosshans, 2018). All reporters were integrated into the *C. elegans* genome in single copy at the same defined genomic locus.

To assess the extent to which transgenes, and thus promoter activity, could recapitulate endogenous rhythmic gene activity, we compared dynamic changes in abundance of the endogenous transcript and its *gfp* mRNA counterpart within the same worm lysates of synchronized worm populations over time. Specifically, we plated starved L1 larvae on food at 25°C and sampled hourly between 22 – 37 hours after plating (**Figure 1C, Figure S2A**). In each of the eight cases that we examined, chosen to represent a variety of peak phases and amplitudes, we observed rhythmic reporter transcript accumulation. Remarkably, the patterns of the endogenous transcripts and their derived reporters were highly similar, i.e., in all tested cases except one, peak phases and amplitudes were comparable (**Figure 1C, Figure S2A**). (We suspect, but have not examined further, that in the one case where we observe a deviation, *R12*.*E2*.*7p* in Fig S2A, the reporter may lack relevant promoter or intronic enhancer elements.) Furthermore, in the case of *F58H1*.*2*, the reporter RT-qPCR time course recapitulated high-amplitude oscillations despite a much more modest oscillatory signal in the ChIP-seq experiment (**Figure S2B**), further supporting the notion that the differences in amplitudes between ChIP-seq and mRNA-seq probably are of technical nature and do not primarily arise from post-transcriptional regulation of the transcripts.

Taken together, these results reveal that the promoters of a variety of oscillating genes are sufficient to recapitulate endogenous transcript dynamics.

### A targeted screen identifies transcription factors involved in molting

Prompted by the above findings, we sought to identify rhythmically active transcription factors involved in molting. Hence, we performed an RNAi screen targeting 92 transcription factors that exhibit transcript level oscillations according to our previous annotation (Hendriks et al., 2014). Specifically, we screened for aberrant developmental progression or molt execution. To obtain such information, we examined luciferase activity in animals that express a luciferase transgene and that are grown in the presence of D-luciferin (Meeuse et al., 2020; Olmedo et al., 2015). This assay detects lethargus by a drop in luminescence at the level of individual animals, allowing us to quantify durations of molts, intermolts and, as a sum of the two, entire larval stages for several animals per condition. We depleted the transcription factors by feeding animals RNAi-expressing bacteria. To control for differences in larval growth among RNAi conditions unrelated to target protein depletion, we performed the experiment in parallel on RNAi deficient *rde-1(ne219)* mutant animals (Tabara et al., 1999) (**Figure 2A**).

**Figure 2:**
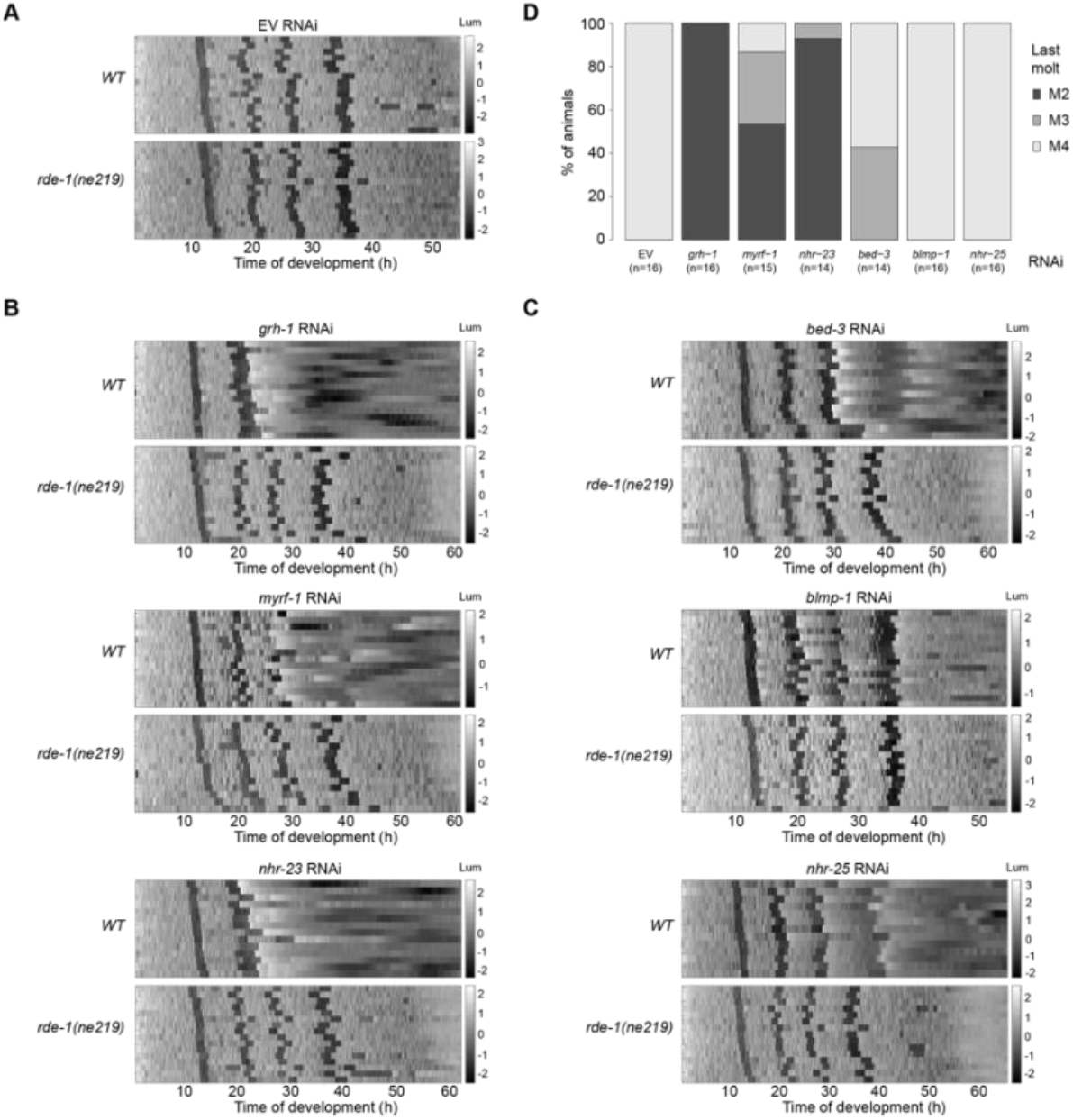
An RNAi screen identifies transcription factors required for normal progression through larval molting cycles. **A**, Heatmap showing trend-corrected luminescence (Lum) of wild-type (WT, strain HW1939, top) and RNAi-deficient (*rde-1(ne219*), strain HW2150, bottom) animals expressing luciferase from the *eft-3* promoter grown on mock (empty vector, EV) RNAi in a temperature-controlled incubator set to 20°C. Each line represents one animal. Hatch is set to t = 0 hours and traces are sorted by entry into the first molt. Darker color corresponds to low luminescence and is associated with the molt. **B, C** Heatmap showing trend-corrected luminescence as in (A), for indicated RNAi conditions causing altered numbers (B) or durations (C) of molts, respectively. **D**, Quantification of the percentage of animals entering specific molts on indicated RNAi conditions. Shown are the last molts observed for animals in each condition; e.g., 100 % of GRH-1 depleted fail to progress beyond M2. See also **Figures S3, S4**

By plotting the luminescence intensities sorted by entry into the first molt in a heatmap (**Figure 2B,C**), we identified six genes whose depletion caused an apparent arrest in development or death following atypical molts (*nhr-23, myrf-1* and *grh-1*; **Figure 2B,D, Figure S3**), or aberrant duration of molts (*bed-3, blmp-1* and *nhr-25*; **Figure 2C,D, Figure S4**).

The nuclear hormone receptors NHR-23 and NHR-25 have previously been shown to function in molting (Frand et al., 2005; Gissendanner et al., 2004; Gissendanner and Sluder, 2000; Johnson et al., 2021; Kostrouchova et al., 1998; Kostrouchova et al., 2001), and we and others have described functions of BLMP-1 in oscillatory gene expression and cuticle formation (Hauser et al., 2021; Sandhu et al., 2021; Stec et al., 2021). Here, we sought to characterize GRH-1, whose orthologues are key regulators of epithelial cell fates and remodeling of epithelial tissues in other animals (Sundararajan et al., 2020), and which had no known role in *C. elegans* post-embryonic development.

### Loss of GRH-1 results in abnormal ecdysis and larval death

Injection of double-stranded RNA targeting *grh-1* into the germline of L4 stage larval animals causes embryonic lethality in the next generation (Venkatesan et al., 2003). Consistent with this finding, we observed an embryonic lethality phenotype in *grh-1(0)* mutant animals (data not shown). To bypass this defect and investigate the role of GRH-1 during larval development, we performed controlled GRH-1 depletion in larvae using the auxin-inducible degradation (AID) system (Zhang et al., 2015). We tagged *grh-1* endogenously with *aid::3xflag* to generate allele *grh-1(xe135)*, and expressed the plant-specific F-box protein TIR1 as a transgene, generating a strain that, for simplicity’s sake, we will refer to as *grh-1::aid* in the following.

When we placed synchronized L1 stage *grh-1::aid* animals on 1 mM auxin-containing plates, we observed fully penetrant lethality: 24 h after plating, 40% of animals had died during an abnormal ecdysis. Surviving animals were variably sized but generally much smaller than wild-type control animals (which were L3/L4), and subsequently also succumbed to failed ecdysis, with no viable larvae left on plate by 38h.

To observe the molting process in greater detail, we used time-lapse DIC imaging to observe L1 animals transferred to an agar pad in a drop of M9 buffer, which allowed them to move. We could readily identify loosened cuticles at the tip of the head in both wild-type and GRH-1-depleted animal (**Figure 3A,B**). Next, animals made spontaneous back-and-forth movements and the pharynx contracted rapidly (**Supplemental Movies 1 & 2**), after which wild-type animals shed the cuticle (ecdysed) (**Figure 3A,B**). By contrast, in GRH-1-depleted animals, the cuticle became even looser and more inflated in the head region (**Figure 3C**). Vesicles appeared in the cavity underneath the loosened cuticle (**Figure 3C**). Finally, the cuticle broke in the head region and the underlying tissue was extruded (**Figure 3C, Supplemental Movie S3**). We conclude that GRH-1 is required for viability at least in part through its role in proper cuticle formation, and that cuticle rupturing during ecdysis provides a likely explanation for the abnormal traces observed for GRH-1-depleted animals in the luciferase assay.

**Figure 3:**
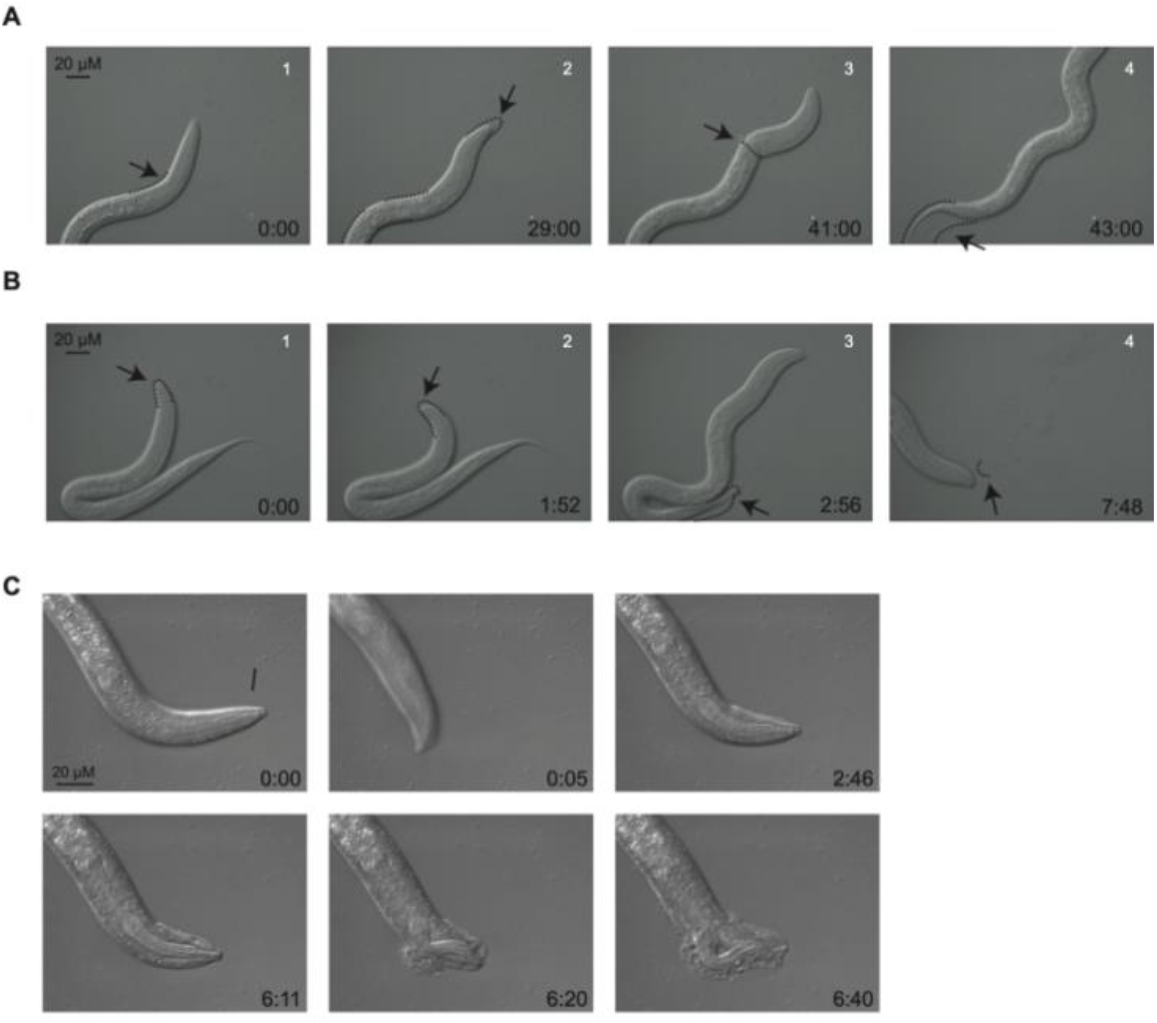
GRH-1 is required for cuticular integrity and normal ecdysis. **A, B**, Image sequence of N2 wild-type animals during M1. Lethargic animals were transferred to an agar pad and observed by DIC at 63x magnification and imaged every 20 s (A) or every 4 s (B) respectively. Selected images with time stamps are shown. Dashed lines indicate cuticle boundaries detached from the body. As reported previously (Singh and Sulston, 1978), two different sequences of events were observed: the pharyngeal lining is removed prior to ecdysis (A), or the pharyngeal lining is expelled after crawling out of the cuticle (B). Arrows indicate specific features of the molt. In (A): 1. Loosened cuticle, 2. Detachment of pharyngeal lining, 3. Crawling out of cuticle, 4. Final crawling out of cuticle. In (B): 1. Loosened cuticle, 2. Back-and-forth movements, 3. Crawling out of cuticle with pharyngeal lining still attached, 4. Pharyngeal lining expelled. **C**, Image sequence of an L1 synchronized *grh-1::aid* animal (HW2418) plated on 250 μM auxin-containing plates, grown at 20°C. A lethargic animal was transferred to an agar-pad containing microscopy slide and images were collected every 1 sec, using DIC, 100x magnification. Selected images of Supplemental Movie 1 are shown. Time stamp (min:sec) is indicated. Arrows indicate phenotypic features: loosening of the cuticle (0:00); back-and-forth movements (0:05); inflation of the cuticle (2:46); vesicles underneath loosened cuticle (6:11); rupturing of the cuticle (6:20, 6:40). See also **Supplemental Movies 1 – 3**

### GRH-1 depletion extends molt durations

We noted an increased duration of molts before GRH-1-depleted animals died (**Figure 2B**, see also Figure **5**, **7B** below). To examine this phenotype further, we exposed *grh-1::aid* animals to varying auxin concentrations to titrate GRH-1 depletion. Although auxin concentrations of ≥400 nM at hatching yielded a fully penetrant M1 phenotype, a dose-dependent decrease in phenotype penetrance occurred below this concentration (**Figure 4A, Figure S5A**). When we quantified the developmental tempo for animals that, at the lowest auxin concentrations tested (53 nM, 79 nM and 119 nM), completed the first three molts, we observed a dose-dependent lengthening in M1, M2 and M3 (**Figure 4B, Figures S5, S6**). This effect increased with each molt. By contrast, little or no change occurred for the duration of the intermolts preceding the lengthened molts (**Figure 4C, Figures S5, S6**). Hence, whereas extensive GRH-1 depletion results in a dysfunctional cuticle and larval death, more modest perturbations change molt durations quantitatively.

**Figure 4:**
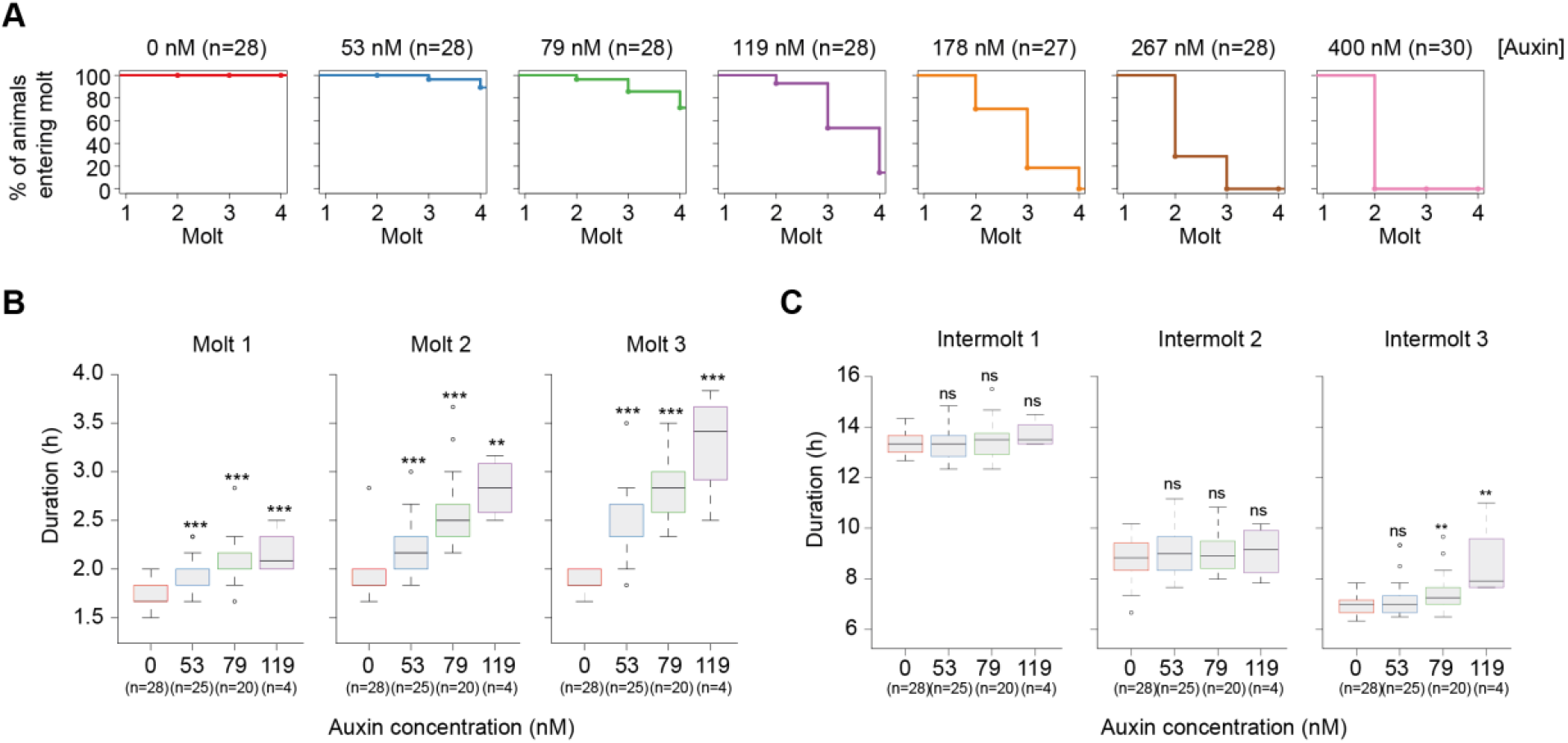
GRH-1 depletion extends molt duration in a dose-dependent manner. **A**, Quantification of the percentage of *grh-1::aid* animals constitutively expressing luciferase (HW2434) that enter each of four molts molt upon hatching into increasing concentrations of auxin as indicated. Molts in single animals grown at 20°C were determined as in **Figure S3**. For concentrations up to 250 μM, similar results as for 400 nM were observed (**Figure S5**). **B**, Boxplot showing the duration of M1, M2 and M3 of animals treated with indicated concentrations of auxin. Animals that failed to develop beyond M3 in (A) were excluded. Significant differences relative to 0 nM auxin are indicated. P-values were determined by Wilcoxon test. ns: not significant, * p<0.05, ** p<0.01, *** p<0.001. See also **Figures SS5, S6**

### GRH-1 is repetitively required for each molt

We wondered if GRH-1 was required for each molt, as we would predict for a component of the core molting machinery. To test this possibility, we utilized timed application of auxin and the luciferase assay. Recapitulating the effect of auxin application to synchronized L1 stage animals grown on a plate, application of auxin at hatch prevented animals from developing beyond M1 (**Figure 5A**). Moreover, when we applied auxin at the beginning of the L2, L3 and L4 stage, respectively, this prevented animals from developing beyond the next respective molt, i.e. M2, M3 and M4 (**Figure 5B – D**). We conclude that GRH-1 is repetitively required during development, for successful completion of each molt. Moreover, defects occur within the stage in which GRH-1 is depleted.

**Figure 5:**
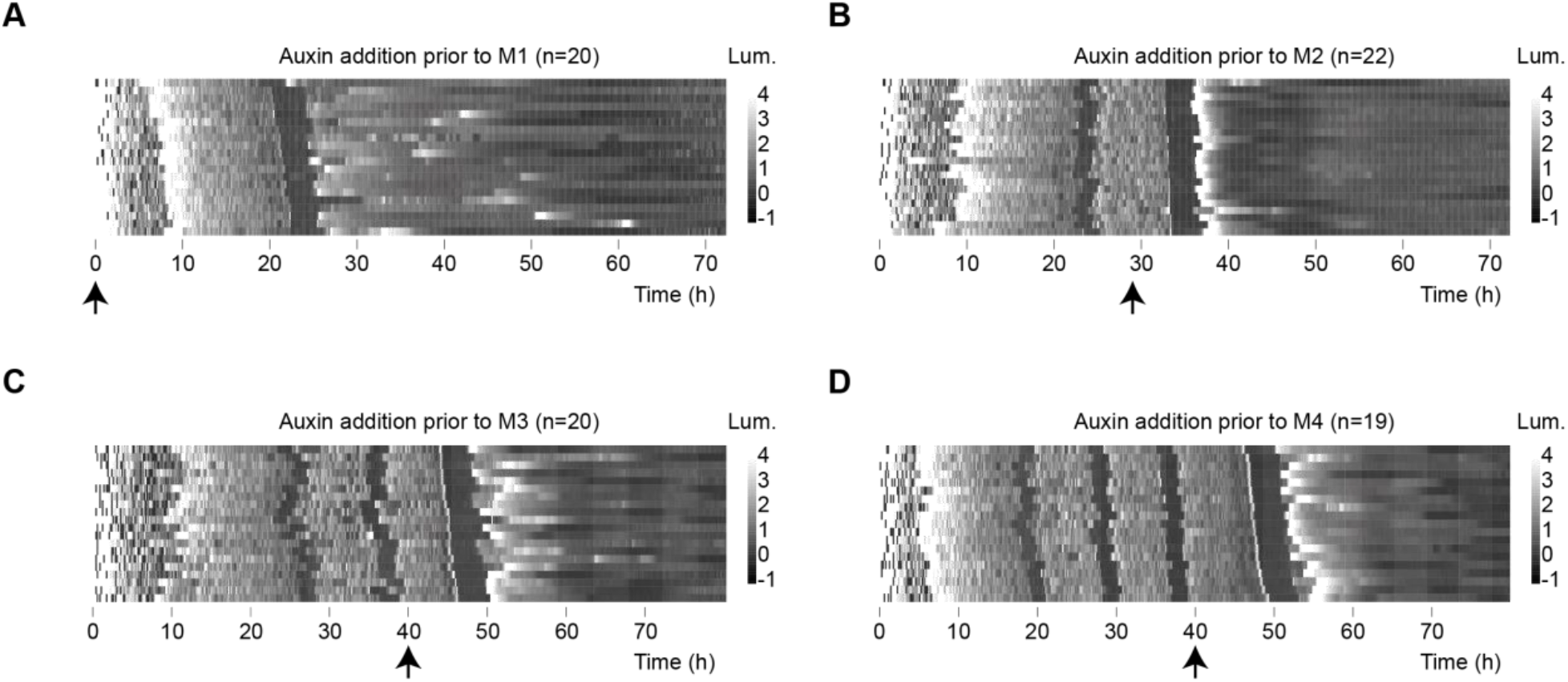
GRH-1 is required for successful completion of each molt. **A-D**, Heatmaps showing trend-corrected luminescence (Lum) of *grh-1::aid* animals constitutively expressing luciferase (HW2434). t = 0h corresponds to time of plating embryos, which subsequently hatch at different times. Arrow indicates time point when 250 μM auxin was added, i.e. prior to the first molt (A; M1), M2 (B), M3 (C) or M4 (D) larval stage. Note that for technical convenience in (A), auxin was provided at time of plating. Animals are sorted by entry into M1 (A), M2 (B), M3 (C), M4 (D), respectively.

### GRH-1 protein levels oscillate and peak shortly before molt entry

Since GRH-1 is required repetitively, for proper execution of each of the four molts, and its mRNA levels oscillate (Hendriks et al., 2014; Meeuse et al., 2020), we wondered whether GRH-1 protein accumulation is also rhythmic. To test this, we examined a GRH-1-GFP fusion protein expressed from the endogenous *grh-1* locus. We observed the first detectable signal in elongating embryos (**Figure S7**), i.e., at the time when oscillatory gene expression initiates (Meeuse et al., 2020). In larvae, GFP signal accumulated in various tissues, including seam cells, vulva precursor cells, non-seam hypodermal cells, rectum cells, socket cells, and pharyngeal cells (**Figure S8**). Finally, in adults, which lack oscillatory gene expression (Meeuse et al., 2020), GFP::GRH-1 levels were greatly diminished or altogether absent in various tissues (**Figure S9**).

To study the dynamics of *grh-1* expression in more detail and relate it to progression through larval development, we performed time-lapse microscopy of single animals grown in micro-chambers, acquiring fluorescence and bright-field images in parallel and with high temporal resolution as described previously (Meeuse et al., 2020). We observed robust rhythmic GRH-1 accumulation with a peak before molt entry (**Figure 6**).

**Figure 6:**
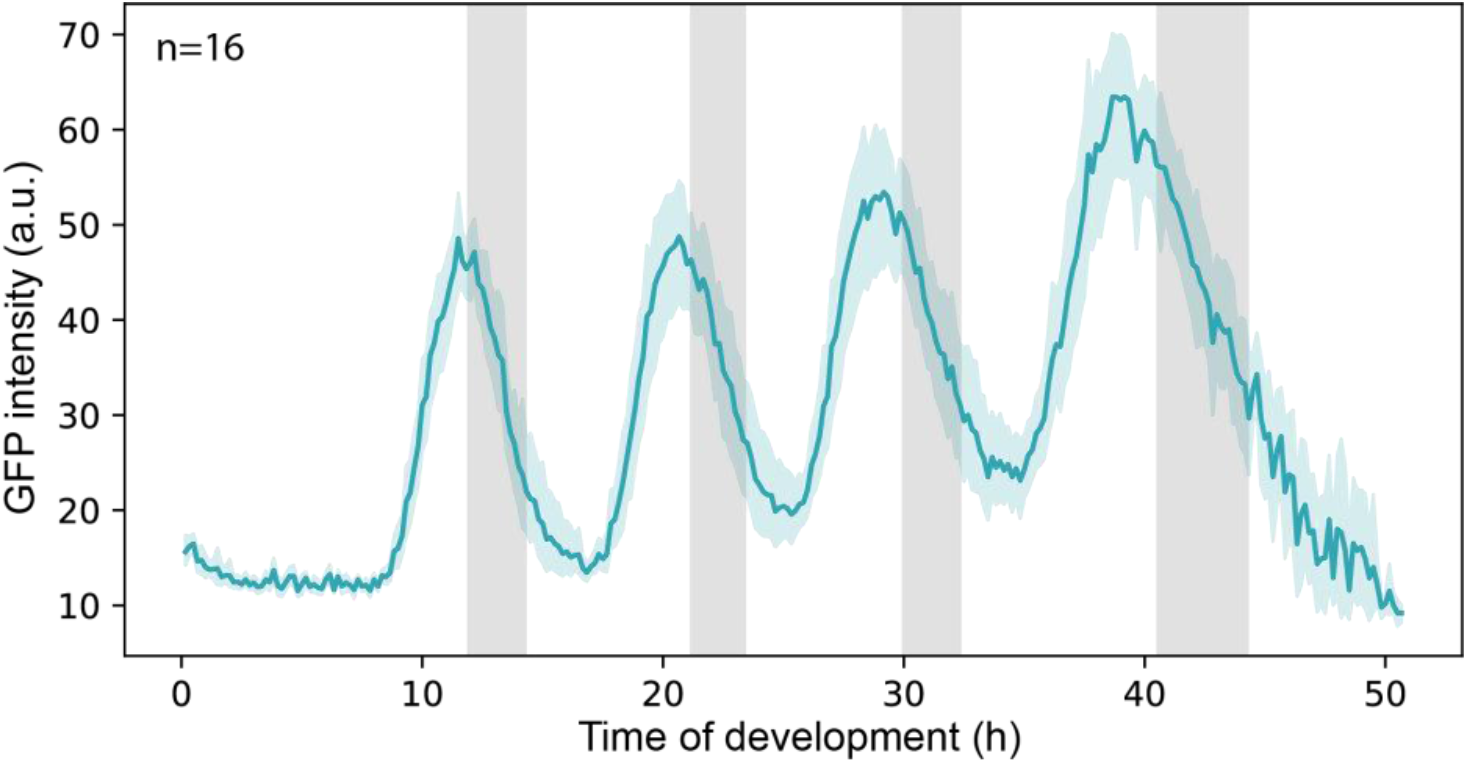
GRH-1 protein accumulates rhythmically before each molt. Time-lapse imaging of animals expressing endogenously tagged *gfp::grh-1*. Average +/-95% confidence interval (cyan shading) are shown; gray boxes indicate average time of molts. See also **Figures S7 – S9**

### Molting requires oscillatory GRH-1 activity

The rhythmic accumulation of GRH-1 suggested that it could also exhibit rhythmic activity, such that it was required only during a certain window of each larval stage to support successful molting. To test this hypothesis, we initiated GRH-1 degradation at variable times in L2 by adding auxin and monitored developmental progression using the luciferase assay. Plotting luminescence traces by the time when the animals entered molt 2 in a heatmap (**Figure 7A,B**) revealed a striking cutoff on the onset of the phenotype: addition of auxin up to 3 hours before the M2 was sufficient for phenotypic onset at M2 exit (**Figure 7B,C**). However, if animals received auxin later than this, i.e., within 3 hours from M2 entry or during M2, they progressed through M2 and instead exhibited the phenotype at M3 exit (**Figure 7B,C**). We observed analogous outcomes when auxin was added in L3 (**Figure S10**). This period of auxin resistance is not explained by slow GRH-1 depletion kinetics since auxin depletes GRH-1 effectively within < 1h (**Figure 7D**). Hence, a period of time exists in each larval stage during which GRH-1 is dispensable for molt completion. In other words, and as we had hypothesized, GRH-1 exhibits rhythmic activity during larval development.

**Figure 7:**
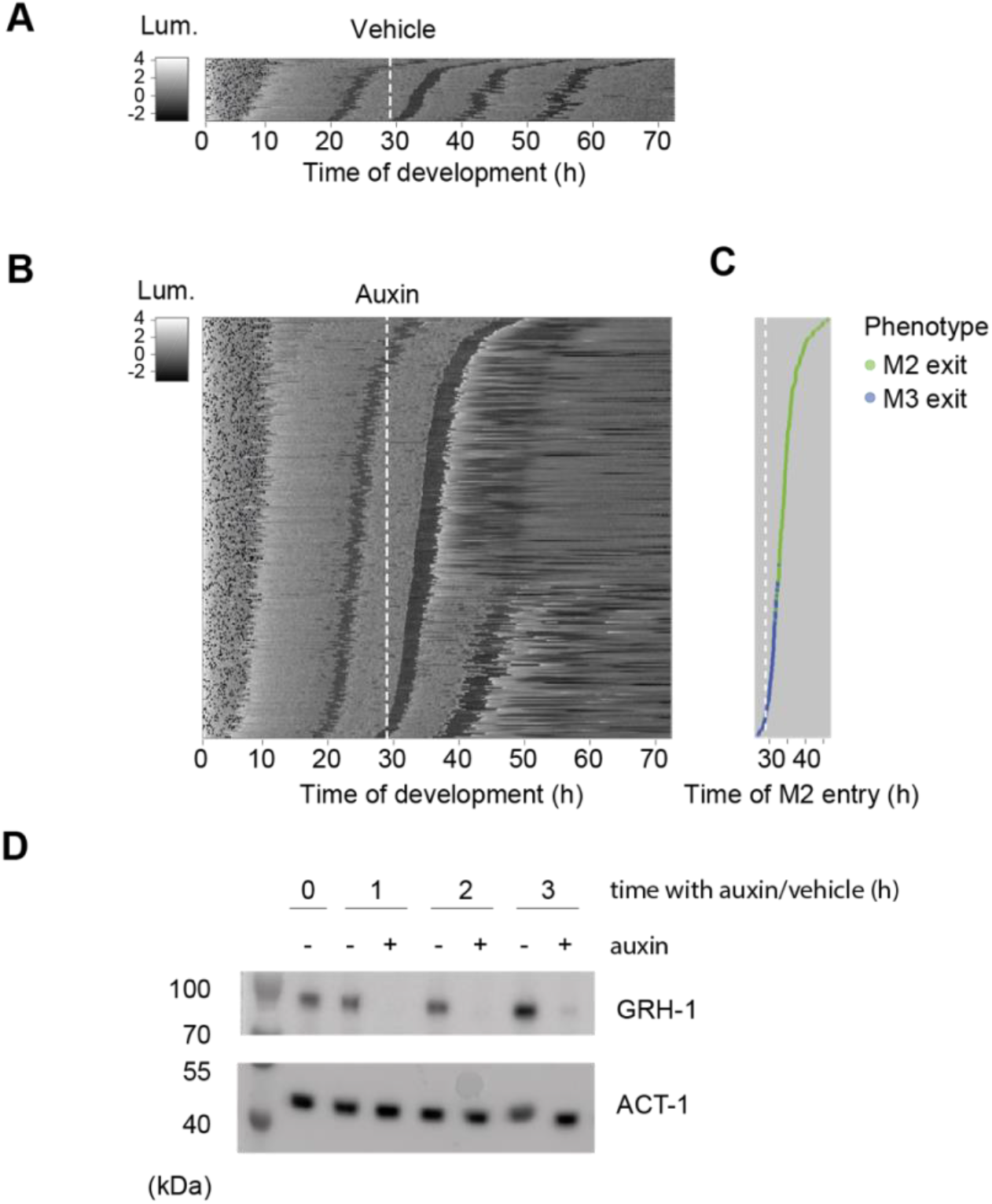
GRH-1 functions prior to the molt. **A**,**B** Heatmap showing trend-correct luminescence (Lum) of *grh-1::aid* animals (HW2434) treated with vehicle (0.25% ethanol (A), or 250 μM auxin (B) at 29 hours after plating (white dashed line). Black intensities reflect low luminescence during lethargus (molt). Embryos of various stages were plated to obtain an asynchronously hatching population. Luminescence traces are sorted by entry into molt 2 (M2) such that traces of early hatched animals are at the bottom and those of late hatched animals are at the top. **C**, Plot of phenotype onset over time of auxin application relative to entry into molt 2 (M2 entry). Dots represent individual animals from (B), colored according to whether the last observed molt in the luminescence trace was M2 (M2 exit phenotype; green) or M3 (M3 exit phenotype; blue). **D**, Western blot revealing rapid GRH-1 depletion in the *grh-1::aid* strain (HW2434) upon addition of 250 μM auxin. A synchronized culture of animals was grown in liquid at 20°C. After 21 h (denoted t=0 h in the figure), the culture was split in two and either auxin or vehicle were added as indicated. Cultures were sampled hourly and protein lysates probed by Western blotting using anti-FLAG and anti-actin antibodies as indicated. See also **Figure S10**.

## Discussion

Although molting is a fundamental feature of nematode development, little is known about the GRNs that control it. This is particularly true of the clock-type mechanisms thought to facilitate the regular occurrence of molts and synchronization of the various processes that they encompass. Here, we have demonstrated that rhythmic transcription drives oscillatory accumulation of a large fraction of genes. We devised a screen that identified six transcription factors important for molting and characterized the function of GRH-1. Our work provides a basis for elucidating the GRNs that support molting timing and execution timing and, potentially, oscillatory gene expression.

### Rhythmic transcription generates oscillating mRNA levels

Our previous work demonstrating rhythmic intronic RNA accumulation (Hendriks et al., 2014) implicated rhythmic transcription in the *C. elegans* larval oscillator. Time-resolved RNAPII ChIP seq and reporter gene assays presented here now provide direct experimental support of this idea: Globally, ChIP-seq and mRNA-seq experiments yielded highly similar patterns, and several promoter fusion transgenes recapitulated the rhythmic accumulation of endogenous transcripts both qualitatively, as also noted in other instances (Frand et al., 2005; Hao et al., 2006), and quantitatively at the level of phases and amplitudes.

Although our results demonstrate a major function of rhythmic RNA polymerase II recruitment in oscillating mRNA accumulation, we emphasize that, at least in specific cases, post-transcriptional regulatory mechanisms may serve to generate, shape or damp oscillations, e.g. through miRNA-mediated repression (Kim et al., 2013). The data we have presented here may allow identification of such instances in the future, although we caution that the apparently lower dynamic range of ChIP-seq poses a technical challenge.

### Identification of six transcription factors important for proper execution of molting cycles

Prompted by the finding that oscillatory mRNA accumulation relies on rhythmic transcription, we decided to screen for transcription factors potentially involved in the process by identifying factors with altered timing or reduced numbers of molts. In principle, the latter phenotype in particular could also be caused by non-specific larval arrest or death, unrelated to functions in developmental timing or molts. It is thus a striking outcome of the screen, and a validation of our approach, that all six hits that we identified are indeed linked to molting, as shown here, in separate work (Hauser et al., 2021), or evidenced by findings from the literature as discussed below. We propose that this reflects the pre-selection of our candidates as rhythmically expressed transcription factors.

Among the hits, we identified NHR-23/RORA, arguably the best characterized molting transcription factor, whose depletion was previously shown to cause a failure of ecdysis and larval arrest (Frand et al., 2005; Gissendanner et al., 2004; Johnson et al., 2021; Kostrouchova et al., 1998; Kostrouchova et al., 2001; Kouns et al., 2011) and which has a role in regulating the expression of collagens and hedgehog-related genes (Kouns et al., 2011).

The GRNs involving NHR-23’s function in molting are not well understood. NHR-23 is orthologous to DHR3/NR1F1, an ecdysone-controlled fly transcription factor important for metamorphosis (Kostrouchova 1998). In metamorphosis, DHR3 activates FTZ-F1 (Ou and King-Jones, 2013), the orthologue of another one of our screen hits, NHR-25/NR5A. Based on this orthology, and the finding that NHR-25 depletion impairs ecdysis, albeit more weakly and only in later molts than that of NHR-23, it was previously speculated that a regulatory interaction between the two proteins might be conserved in *C. elegans* and contribute to the molting process (Gissendanner et al., 2004; Gissendanner and Sluder, 2000). However, this notion has remained controversial (Kostrouchova et al., 2001), suggesting that further effort will be needed to understand whether and how NHR-25 – and NHR-23 – function in the molting cycle beyond their immediate regulation of cuticular components and their processing machinery.

Elsewhere, we have reported a detailed characterization of another screen hit, BLMP-1, which we find to be important for both molting timing and oscillatory gene expression (Hauser et al., 2021), possibly through its function as a pioneer transcription factor (Stec et al., 2021). Consistent with our identification of BED-3 as a screen hit, BLMP-1 was previously reported to promote expression of *bed-3* and partially phenocopy its mutant phenotypes (Yang et al., 2015). Finally, MYRF-1, also known as PQN-47, is required for ecdysis (Meng et al., 2017; Russel et al., 2011).

### The Grainyhead transcription factor GRH-1 plays an important role in molting

Here, we have focused on characterizing GRH-1. Previously the least studied factor among the six hits, it is a member of the Grainyhead/LSF1 protein family that controls epithelial cell fates across animals (Sundararajan et al., 2020). We showed that it is repetively required for molting, during a specific window of each larval stage, before late intermolt. This window is consistent with, and presumably determined by, the rhythmic accumulation of GRH-1 protein, whose levels peak shortly before molt entry.

We show that the failure to complete development observed in the screen is due to a cuticle and ecdysis defect: the onset of ecdysis is delayed and the newly formed cuticle ruptures during ecdysis, particularly in the larval head region. Consistent with the view that generation of the new cuticle is defective, rupturing happens specifically after the onset of ecdysis, with tissue extrusion occurring after the old (outer) cuticle has already visibly detached from the worm body. Indeed, the notion of impaired cuticle biogenesis is also consistent with cuticle defects in GRH-1-depleted *C. elegans* embryos, which do not undergo ecdysis (Venkatesan et al., 2003), although it remains possible that inappropriate proteolytic activities during ecdysis might additionally damage the newly formed cuticle.

Soft, thin and granular cuticles, prone to rupturing, are also a hallmark of *Drosophila* Grainyhead mutant animals (Bray and Kafatos, 1991; Nüsslein-Volhard et al., 1984). In mammals, *Grhl1* and *Grhl3* are dynamically expressed in the epidermis (Joost et al., 2020; Joost et al., 2016) and control epidermal differentiation (Yu et al., 2006), wound healing and eye lid closure (Boglev et al., 2011). Hence, our findings agree with a fundamental, evolutionarily conserved role of Grainyhead proteins in epidermis and ECM development and remodeling. It will thus be interesting to determine, in future work, the direct transcriptional outputs of GRH-1 and the GRNs that enable its dynamic expression. Such work may then also help to understand whether the delayed onset of ecdysis observed with sub-lethal GRH-1 depletion reflects a genuine function of GRH-1 in timing this process or an indirect outcome of perturbed cuticle formation.

Finally, we note that the orthologues of additional screen hits function in development and regeneration of mammalian skin and its appendages, notably hair. We are particularly intrigued by the apparent parallels between *C. elegans* molting and the mammalian hair follicle cycle. This process of rhythmic homeostatic skin regeneration is controlled by a poorly characterized “hair follicle clock”, modulated by circadian rhythms (Paus and Foitzik, 2004; Plikus et al., 2015). Mutations in the mouse orthologues of *blmp-1* and *nhr-23, Prdm1* and *Rora*, respectively, impair the proper execution of hair follicle cycles (Magnusdottir et al., 2007; Steinmayr et al., 1998; Telerman et al., 2017), while *grhl1* ablation causes hair coat defects related to hair anchoring and potentially other processes (Wilanowski et al., 2008).

In summary, we propose that in this study we have identified and characterized a number of factors linked to a molting cycle oscillator that may have a functional correspondence in other animals, and that their further dissection provides a path to a molecular-mechanistic and systems understanding of gene expression oscillations in *C. elegans*. Additionally, we propose that *C. elegans* molting could serve as a powerful, genetically accessible, model of animal skin development and regeneration.

## Methods

### *C. elegans* strains

The Bristol N2 isolate was used as the wild-type reference. We generated or used the following additional strains.

*rde-1(ne219) V* (Tabara et al., 1999)

HW1360: EG6699, *xeSi131[F58H1*.*2p::GFP::H2B::Pest::unc-54 3’, unc-119+] II* (this study)

HW1361: EG6699, *xeSi132[R12E2*.*7p::GFP::H2B::Pest::unc-54 3’, unc-119+] II* (this study)

HW1370: EG6699; *xeSi136[F11E6*.*3p::GFP-H2B-Pest::unc-54 3’UTR; unc-119 +] II* (Meeuse et al., 2020)

HW1371: EG6699; *xeSi137[F33D4*.*6p::GFP-H2B-Pest::unc-54 3’UTR; unc-119 +] II* (this study)

HW1372: EG6699; *xeSi138[C05C10*.*3p::GFP-H2B-Pest::unc-54 3’UTR; unc-119 +] II* (this study)

HW1431: EG6699, *xeSi160[daf-6Δ4p::GFP::H2B::Pest::unc-54 3’UTR; unc-119 +] II* (this study)

HW1939: *EG6699, xeSi296 [eft-3p::luc::gfp::unc-54 3’UTR, unc-119(+)] II* (Meeuse et al., 2020)

HW1949: *EG8080, xeSi301 [eft-3p::luc::gfp::unc-54 3’UTR, unc-119(+)] III* (this study)

HW1984: *ieSi57 [eft-3p::TIR1::mRuby::unc-54 3’UTR, cb-unc-119(+)] II; EG8080, xeSi301 [eft-3p::luc::gfp::unc-54 3’UTR, unc-119(+)] III* (this study)

HW2079: *EG8080, xeSi376 [eft-3p::TIR1::mRuby::unc-54 3’UTR, cb-unc-119(+)] III* (this study)

HW2150: *EG6699, xeSi296 [eft-3p::luc::gfp::unc-54 3’UTR, unc-119(+)] II; rde-1(ne219) V* (this study)

CA1200: *ieSi57 [eft-3p::TIR1::mRuby::unc-54 3’UTR, cb-unc-119(+)] II* (Zhang et al., 2015)

HW2418: *grh-1(xe135(grh-1::aid::3xflag)) I; EG8080, xeSi376 [eft-3p::TIR1::mRuby::unc-54 3’UTR, cb-unc-119(+)] III* (this study)

HW2434: *grh-1(xe135(grh-1::aid::3xflag)) I; EG6699, xeSi296 [eft-3p::luc::gfp::unc-54 3’UTR, unc-119(+)] II; EG8080, xeSi376 [eft-3p::TIR1::mRuby::unc-54 3’UTR, cb-unc-119(+)] III* (this study)

HW2493: *grh-1(xe146) I; hT2* (this study)

HW2526: *EG6699, xeSi440[dpy-9p::GFP::H2B::Pest::unc-54 3’UTR; unc-119 +] II* (Meeuse et al., 2020)

HW2533: *EG6699; xeSi442[col-10p::GFP::H2B::Pest::unc-54 3’UTR; unc-119 +] II* (this study)

HW2603: *grh-1(syb616(grh-1::gfp::3xflag)) I* (PHX616, three times backcrossed) (this study; custom-made by SunyBiotech)

### Generation of transgenic animals

Endogenous *aid::3xflag* tagging *grh-1* and generation of *grh-1(0)* mutants by CRISPR/Cas9-mediated editing was performed using the previously published *dpy-10(cn64)* co-conversion protocol (Arribere et al., 2014). For the sgRNA plasmid, we inserted the sgRNA sequence (5’ agaggtttactctcatgagt 3’) into NotI-digested pIK198 (Katic et al., 2015) by Gibson assembly (Gibson et al., 2009).

For *aid::3xflag* tagging, we used hybridized MM116 (5’ AATTGCAAATCTAAATGTTTagaggtttactctcatgagtGTTTAAGAGCTATGCTGGAA 3’) and MM117 (5’ TTCCAGCATAGCTCTTAAACactcatgagagtaaacctctAAACATTTAGATTTGCAATT 3’). A*n aid::linker::3xflag*-linker cassette was synthesized as gBlocks® Gene Fragments (Integrated DNA Technologies) with 50 bp homology arms to the *grh-1* locus before the stop codon: 5’ CCACGTTAATCGAGGTGGCTCCCACCAATCCAAACTCGTATTCCAACTCAATGCCTAAAGATCCAGCCAAACCTCCGGCCAAGGCACAAGTTGTGGGATGGCCACCGGTGAGATCATACCGGAAGAACGTGATGGTTTCCTGCCAAAAATCAAGCGGTGGCCCGGAGGCGGCGGCGTTCGTGAAGAGTACCTCAGGCGGCTCGGGTGGTACTGGCGGCAGCGACTACAAAGACCATGACGGTGATTATAAAGATCATGACATCGATTACAAGGATGACGATGACAAGAGTACTAGCGGTGGCAGTGGAGGTACCGGCGGAAGCTGAGAGTAAACCTCTTTAGGTTCTTGTCTTAATTCTCTTAAAGGAGGACT 3’. Wildtype animals were injected with 10 ng/μl gBlock, 100 ng/μl sgRNA plasmid, 20 ng/μl AF-ZF-827 (Arribere et al., 2014), 50 ng/μl pIK155 (Katic et al., 2015) and 100 ng/μl pIK208 (Katic et al., 2015). Genome editing was confirmed by sequencing.

For *grh-1(0)*, we used hybrized MM116 and MM117, and hybrized MM118 (5’ AATTGCAAATCTAAATGTTTGCGAAAATGCAAACATTAGAGTTTAAGAGCTATGCTGGAA 3’) and MM119 (5’ TTCCAGCATAGCTCTTAAACTCTAATGTTTGCATTTTCGCAAACATTTAGATTTGCAATT 3’) to delete the coding region of *grh-1* and generate the *grh-1(xe146)* allele. Wildtype animals were injected with 100 ng/μl of each sgRNA plasmids, 19.2 ng/μl AF-ZF-827 (Arribere et al., 2014), 47.9 ng/μl pIK155 (Katic et al., 2015) and 95.8 ng/μl pIK208 (Katic et al., 2015). Genome editing was confirmed by sequencing. *grh-1(xe146)* homozygous animals could not be maintained and the *grh-1(xe146)* allele was balanced with hT2.

### Transgenic reporter strain generation

GFP reporters were cloned by Gibson assembly (Gibson et al., 2009). Promoters sequences were amplified from genomic DNA using the primers listed below (overhangs indicated in bold) and inserted into *Nhe*1-digested pYPH0.14 as previously described, (Meeuse et al., 2020). Transgenic animals were obtained by single copy-integration of the transgene into the ttTi5605 locus (MosSCI site) on chromosome II into EG6699 animals, using the published MosSCI protocol (Frøkjær-Jensen et al., 2012).

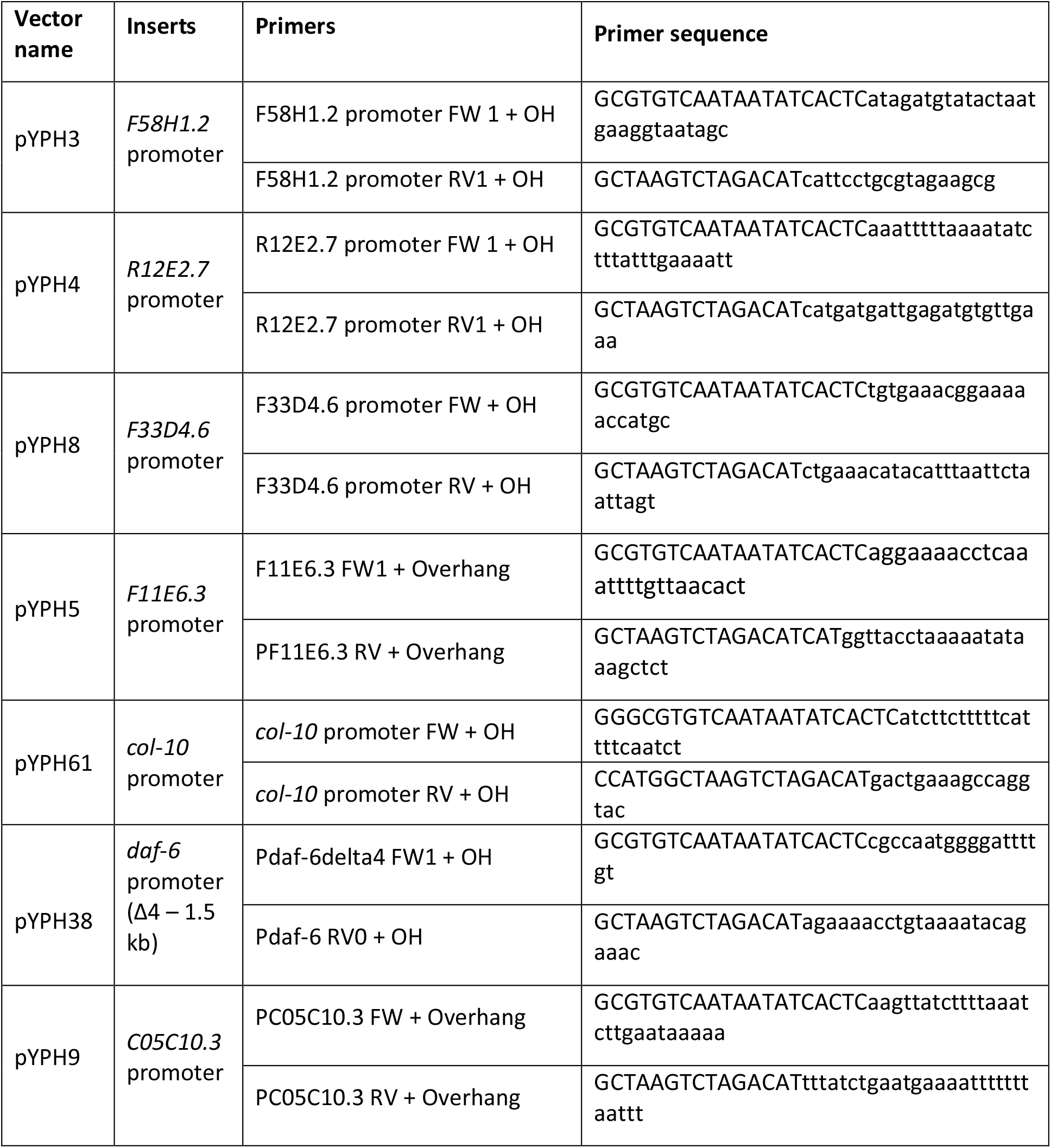

### RNA polymerase II ChIP-sequencing

For RNA polymerase II ChIP-sequencing, synchronized L1 wild-type worms were grown at 25°C. Worms were collected hourly from 22 hours (90,000 worms) until 33 hours (46,000 worms) developmental time. RNA polymerase II ChIP was performed as previously described (Miki et al., 2017). In short, worms were incubated in M9 with 2% formaldehyde for 30 minutes at room temperature with gentle agitation to allow protein-DNA crosslinking. Worms were lysed with beads using the FastPrep-24 5G machine (MP Biomedicals, settings: 8 m/sec, 30 sec on, 90 sec off, 5 cycles). Lysates were sonicated using the Bioruptor Plus Sonication system (Diagenode, settings: 30 sec on, 30 sec off, 20 cycles). 250 μg sonicated chromatin was incubated with 10 μg mouse anti-RNA polymerase II CTD antibody (8WG16, Abcam) at 4°C for 2 hours with gentle agitation and subsequently with 45 μL Dynabeads Protein G (Thermo Fisher Scientific) at 4°C overnight with gentle agitation. Eluate was treated with 0.13 μg/μl RNase and 1 μg/μl Proteinase K. ChIP-seq libraries were prepared using NEBNext Ultra DNA Library Prep Kit for Illumina (New England Biolabs) and sequenced using the HiSeq 50 cycle single-end reads protocol on the HiSeq 2500 system.

Sequencing reads were aligned to the ce10 *C. elegans* genome using the qAlign function (default parameters) from the QuasR package (Gaidatzis et al., 2015) in R. ChIP-seq counts within 1-kb windows, i.e. -500 bp to +500 bp around the annotated TSS (using WS220/ce10 annotations), were scaled by total mapped library size per sample and log_2_-transformed after adding a pseudocount of 8. Genes with a mean scaled TSS window count of less than 8 across all samples were excluded. Log_2_-transformed counts were then quantile-normalized using the normalize.quantiles function from the preprocessCore library (Bolstad, 2021; Bolstad et al., 2003) in R. Finally, quantile-normalized values were row-centered.

### RNA-sequencing for PolII ChIP-seq

For total RNA sequencing, worms were collected hourly from 22 hrs (10,000 worms) until 33 hrs (5,000 worms per time point) after plating from the same plates as for ChIP sequencing. RNA was extracted in Tri Reagent and DNase treated as described previously (Hendriks et al., 2014). Total RNA-seq libraries were prepared using Total RNA-seq ScriptSeq Library Prep Kit for Illumina (New England Biolabs) and sequenced using the HiSeq 50 cycle single-end reads protocol on the HiSeq 2500 system. RNA-seq data were mapped to the *C. elegans* genome (ce10) using the qAlign function (splicedAlignment=TRUE, Rbowtie aligner version 1.16.0) from the QuasR package in R (Gaidatzis et al., 2015) (version 1.16.0). Exonic expression was quantified using qCount function from the QuasR package in R and the exon annotation of the ce10 assembly (version WS220). Counts were scaled by total mapped library size for each sample. A pseudocount of 8 was added and counts were log2-transformed. Lowly expressed genes were excluded (maximum log2-transformed exonic expression - (log2(gene width) - mean(log2(gene width))) ≤ 6). Of the previously annotated ‘high-confidence-oscillating’ genes (n=3,739) (Meeuse et al., 2019), 2,106 genes were sufficiently expressed on the exonic level at the chosen sequencing depth. Oscillating genes were sorted by phase and mean-normalized expression was plotted in heatmaps.

### RT-qPCR reporters

Gravid adult worms were bleached to obtain eggs which were incubated in M9 buffer overnight (12 to 16 hours) on a rotating wheel. After incubation, hatched worms will be synchronized in L1 arrest due to starvation. The synchronized L1 population was plated onto agar plates with food (*E. coli, OP50*) to initiate synchronous larval development. The concentration of worms per plate can vary between 1,000 and 4,000 worms per plate. In total, 2,000 – 8,000 worms were sampled each time point with fewer worms for the last time points.

Worms were collected hourly between 22 and 37 hours at 25°C (for *gfp* reporter data) after plating synchronized L1. Worms were washed off the plate(s) and washed 3 times in M9 buffer. After washing, 1ml Tri Reagent (MRC) was added. Samples were frozen in liquid nitrogen and stored overnight at -80°C. Conventional RNA isolation using phenol chloroform extraction (adapted from (Bethke et al., 2009) was used to extract RNA which was then diluted to the same concentration for each sample and used as input for the Promega Protocol: “ImProm-II™ Reverse Transcription System” to convert RNA to cDNA. The resulting cDNA was diluted 1:1000 to quantify actin transcript levels and 1:20 for endogenous transcripts. qPCR was then performed on a Step one Realtime PCR machine using primer pairs of which one was exon-exon spanning to detect mature mRNA levels of endogenous genes corresponding to the reporters and the *gfp* transcript.

### RT-qPCR primers

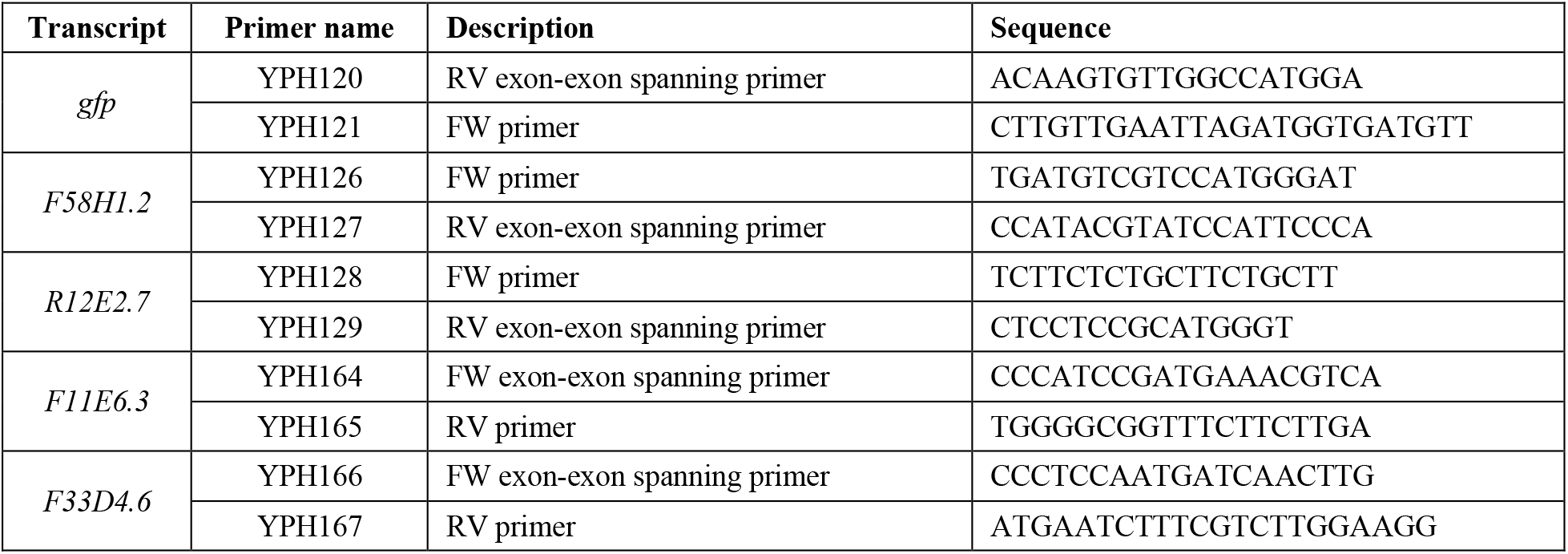
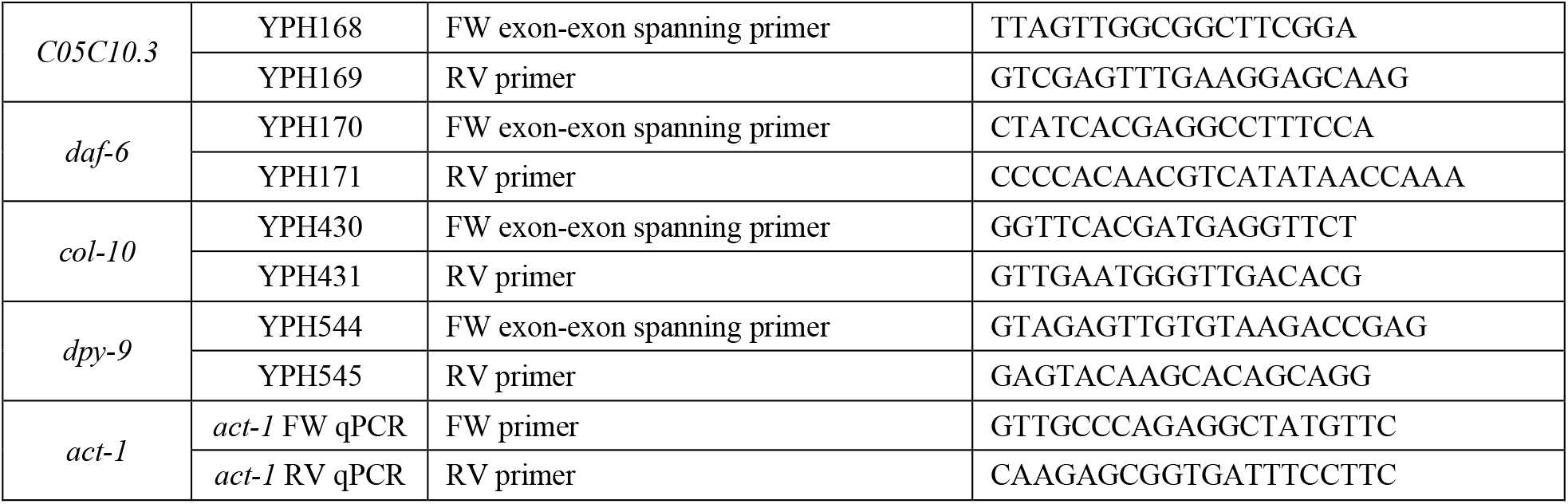

### RT-qPCR analysis

From two technical replicates, the mean actin Ct values were subtracted from the mean target Ct values, to obtain a relative quantification, represented by delta Ct (dCt). To obtain the mean normalized mRNA levels, the dCt mean of the time series was subtracted from from each time point value first and then multiplied by -1. These values were then plotted to compare endogenous versus gfp mRNA levels.

### Luciferase assays

Luciferase assays were performed as described before (Meeuse et al., 2020). In short, single embryos, expressing luciferase from a constitutive and ubiquitous promoter (transgene *xeSi296*) were placed in a 384-well plate (Berthold Technologies, 32505) by pipetting, and left to develop until adulthood in 90 μl S-Basal medium containing *E. coli* OP50 (OD_600_ = 0.9) and 100 μM Firefly D-Luciferin (p.j.k., 102111). Luminescence was measured using a luminometer (Berthold Technologies, Centro XS3 LB 960) every 10 minutes for 0.5 sec for 72 hours in a temperature controlled incubator set to 20 degrees.

For auxin experiments, a 400x stock solution of 3-indoleacetic acid (auxin, Sigma Aldrich, I2886) in 100% ethanol was prepared. The stock solution was diluted 400-fold in the culture medium at the start of each experiment or at specific time points and in concentrations as indicated.

Luminescence data was analyzed using an automated algorithm to detect the hatch and the molts in MATLAB with the possibility to annotate molts manually, as described before (Meeuse et al., 2020). Completion of molts was scored by the presence of a drop in luminescence, followed by a period of stable and low luminescence and subsequent rise in luminescence.

### RNAi screen

To knock-down 92 ‘oscillating’ transcription factors, we used the RNAi feeding method. *E. coli* HT115 bacteria carrying RNAi plasmids were obtained from either of the two libraries (Ahringer library (Fraser et al., 2000; Kamath et al., 2003), Vidal library (Rual et al., 2004)) or cloned (see generation of RNAi vectors).

Luciferase assays were performed as described above with the following adaptations: RNAi bacteria were grown in 5 ml auto-induction medium (2 mM MgSO_4_, 3.3 g/l (NH_4_)2SO_4_, 6.8 g/l KH_2_PO_4_, 7.1 g/l Na_2_HPO_4_, 5 g/l glycerol, 0.5 g/l glucose, 2 g/l α-lactose, 100 μg/ml Amp in ZY medium (10 g/l tryptone, 5 g/l yeast extract)) at 37 °C. Bacteria were diluted in S-Basal medium (OD600 = 0.45), with 100 μM Firefly D-luciferin (p.j.k., 102111) and 100 μg/ml Ampicillin.

We used HW1939 animals that express the *xeSi296* transgene. As a control strain, we used HW2150 animals expressing *xeSi296* in an *rde-1(ne219)* (Tabara et al., 1999) background, which are RNAi deficient. For each RNAi condition we used 2 adjacent columns in the 384-wells plate, i.e. 32 wells with 90 μl culture medium each. To avoid plate effects, the first 8 wells of the first column and the last 8 wells of the second column of the same RNAi condition were filled with an HW1939 animal and the remaining wells with an HW2150 animal.

To identify mutants, we inspected the heatmaps with trend-corrected luminescence (Olmedo et al., 2015) for aberrant duration or number of molts and intermolts.

### Generation of RNAi vectors

For clones that were not available in the Ahringer or Vidal libraries, cDNA or genomic DNA was PCR amplified using the following primers:

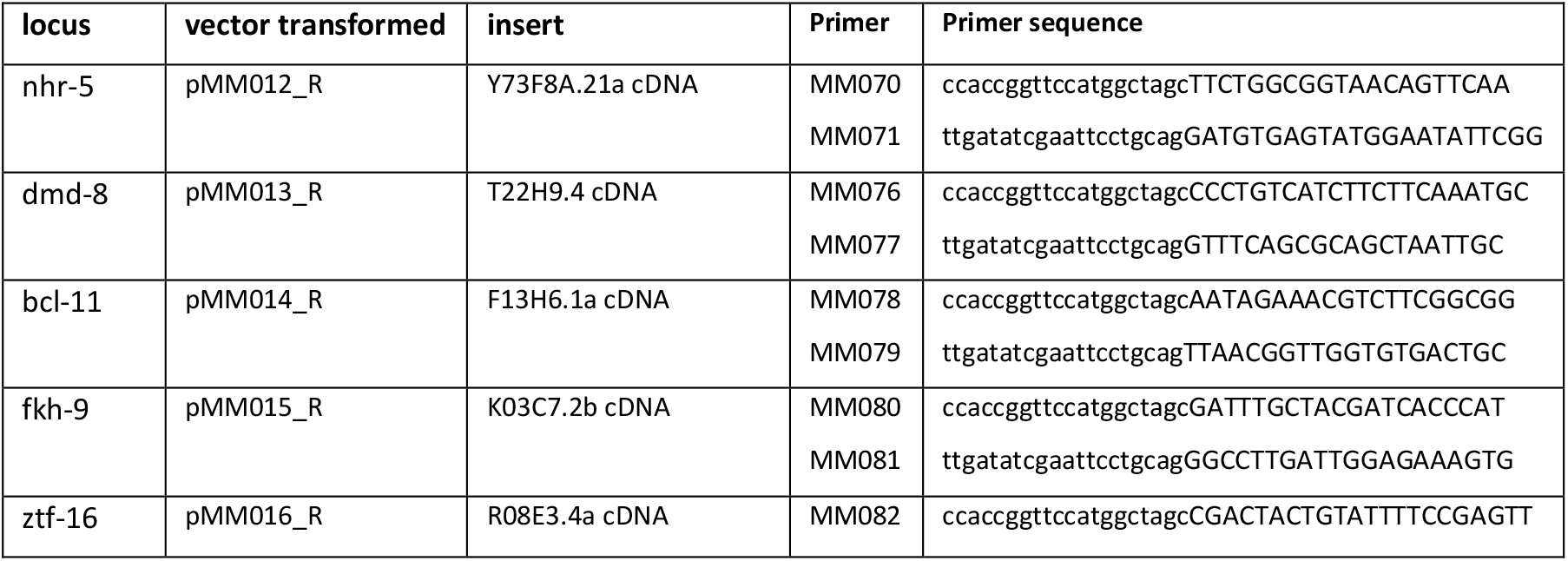
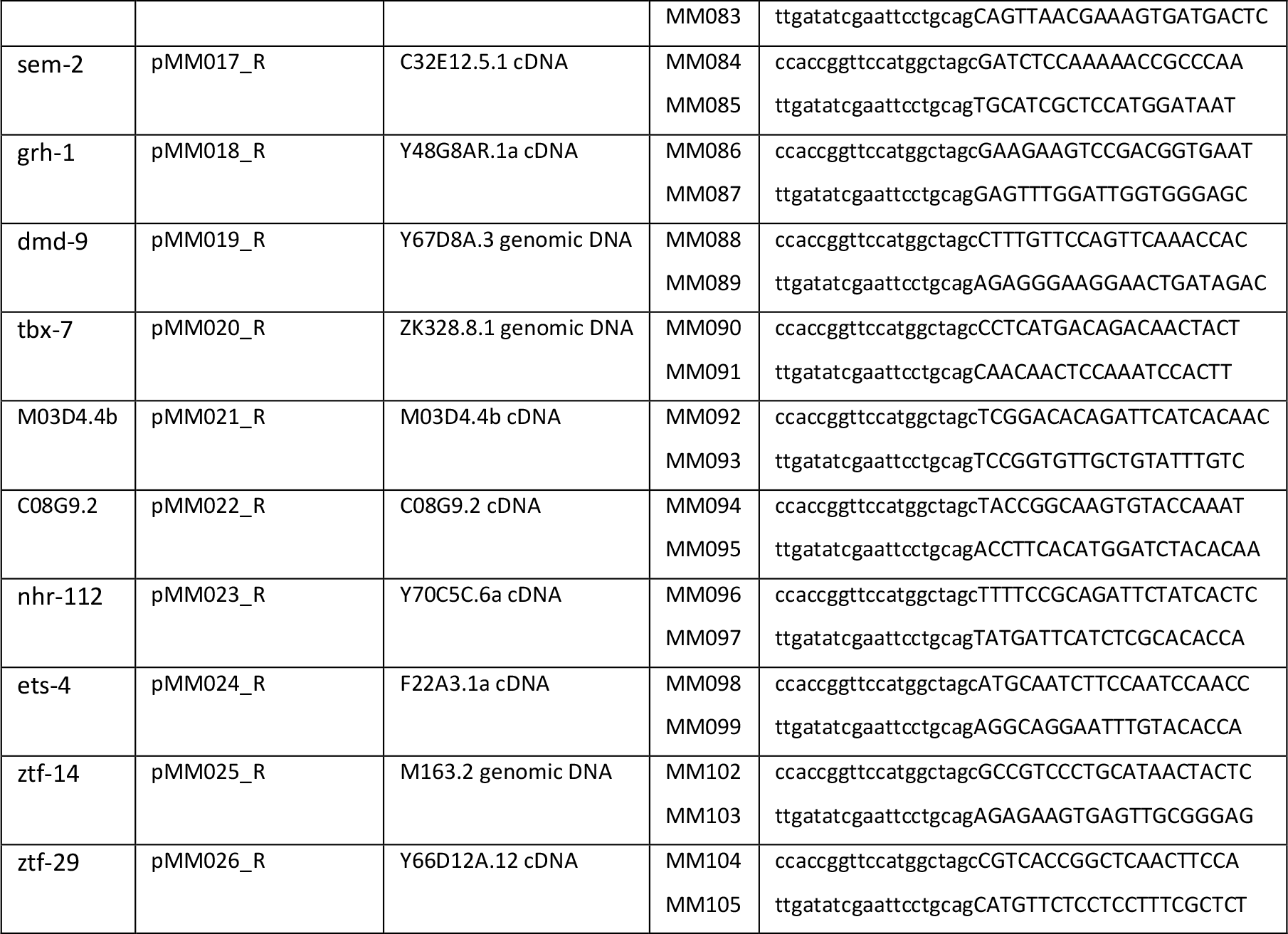

PCR fragments were cloned into the RNAi feeding Pml1 and Sma1 digested L4440 vector (L4440 was a gift from Andrew Fire (Addgene plasmid # 1654 ; http://n2t.net/addgene:1654 ; RRID:Addgene_1654)) using Gibson assembly (Gibson et al., 2009)) and transformed into *E. coli* HT115 bacteria.

### Phenotype imaging

For imaging phenotypes, HW2418 worms were mounted on a 2% (w/v) agarose pad with a drop of M9 buffer (42 mM Na_2_HPO_4_, 22 mM KH_2_PO_4_, 86 mM NaCl, 1 mM MgSO_4_). *grh-1::aid* animals were imaged on an Axio Imager Z1 (Zeiss) microscope. We acquired Differential Interference Contrast (DIC) images using a 100x/1.4 oil immersion objective and a TL Halogen Lamp (3.00 Volt, 900 ms exposure). Images (1388×1040 pixels, 142.1 μm x 106.48 μm pixel size, 12 Bit) were acquired every second from the moment that the cuticle became loose around the tip of the head until after the worm burst through the head, which took roughly 5 to 10 minutes. N2 animals were imaged on an Axio Imager Z2 (Zeiss) microscope. We acquired Differential Interference Contrast (DIC) images using a 63x/1.4 oil immersion objective) and a TL Vis-LED Lamp (5.74 Volt, 17 ms (**Figure 3A**) or 19 ms (**Figure 3B**) exposure). Images (1388×1040 pixels, 225.56 μm x 169.01 μm pixel size, 12 Bit) were acquired at 20 seconds ((**Figure 3A**) or 4 seconds (**Figure 3B**) intervals from the moment that the cuticle became loose around the tip of the head until the cuticle was shed.

### Single worm imaging

To investigate the temporal expression pattern of endogenously tagged *grh-1::gfp*, animals of the strain HW2603 (*grh-1(syb616(grh-1::gfp::3xflag)) I*) were observed by time lapse imaging using single worm imaging as previously described (Meeuse et al., 2020) with slight modifications. Briefly, an array of microchambers (Bringmann, 2011; Turek et al., 2015) was made of 4.5% agarose in S-Basal medium (Stiernagle, 2005). OP50 bacteria were grown on agar plates, scraped off, and transferred to the chambers. Single eggs were placed on the chambers and flipped into a glass coverslip surrounded by a silicone insulator. Low melting agarose (3.5%) was used to seal the edges of the array, which was subsequently mounted on a glass slide for imaging. We imaged animals using a 2x sCMOS camera model (T2) on an Axio Imager M2 (upright microsocope) CSU_W1 Yokogawa spinning disk microscope with a 20x air objective (NA=0.8). The 488 nm laser was set to 70%, with 10 ms exposure and a binning of 2. The brightfield and fluorescent images were taken in parallel using a motorized z-drive with a 2-μm step size and 23 images per z-stack for a total duration of 60h at 10 min time intervals in a ∼21°C room.

### Single worm imaging data analysis

Brightfield images were segmented using a Convolutional Neural Network (CNN v2) (to be communicated elsewhere). Using a default threshold of 127 on the segmentation probability, GFP intensities were quantified on the segmented images using a previously published KNIME workflow (Meeuse et al., 2020). In short, worms are straightened and the GFP intensity of the worm is max projected to one pixel line for each time point. Background-subtracted mean GFP intensities are determined from 20%-80% of the anterior-posterior axis for each time point.To annotate molts, each image was visually inspected for molt entry and molt exit by scrolling through z-stack of the individual timepoints. The GFP intensities and lethargus data was plotted together in Python v3.9 using the Seaborn package.

### Western blot

To examine the kinetics of GRH-1-AID-3xFLAG depletion by auxin, L1 synchronized *grh-1(xe135); eft-3p::luc; eft-3p::TIR1* animals (HW2434) were cultured in liquid (S-Basal supplemented with OP50, OD_600_=3, 1,000 animals/ml) at 20°C. After 21 hours, when animals had reached early L2 stage, the culture was sampled, split in two and supplemented with 250 μM auxin or an equivalent amount of vehicle, respectively, followed by hourly sampling. At each time point, 10,000 animals were collected and washed three times with M9 buffer (42 mM Na_2_HPO_4_, 22 mM KH_2_PO_4_, 86 mM NaCl, 1 mM MgSO_4_). Lysates were made by disruption (FastPrep-24, MP Biomedicals, 5 cycles, 25 sec on, 90 sec off), sonication (Biorupter, Diagnode, 10 cycles, 30 sec on, 60 sec off) and subsequent boiling. Proteins were separated by SDS-PAGE and transferred to a PVDF membrane by semi-dry blotting. Antibodies were used at the following dilutions: mouse anti-FLAG-HRP (1:1,000, A8592, Sigma Aldrich), mouse anti-Actin, clone C4 (1:5,000, MAB1501, Millipore), mouse IgG HRP linked (1:7,500, NXA931V, GE Healthcare). We used ECL Western Blotting detection reagent (RPN2232 and RPN2209, GE healthcare) and ImageQuant LAS 4000 chemilumunescence imager (GE Healthcare) for detection.

## Acknowledgements

We thank Iskra Katic and Lan Xu for help in generating transgenic strains, Anca Neagu for support in analyzing *grh-1* mutant phenotypes, Marit van der Does, Markus Rempfler, Benjamin Titze, Jan Eglinger and Laurent Gelman for help with imaging and image analysis, Dimos Gaidatzis for help in analyzing luciferase screening data, Sarah H. Carl for help with RNA Pol II ChIP-seq analysis, and Iskra Katic for comments on the manuscript.

## Author contributions

Y.P.H. performed transcriptional reporter RT-qPCR time courses. K.B. imaged the N2 molt. S.N. performed GRH-1 time-lapse imaging and related data analysis. M.W.M.M performed the remaining experiments and analyzed the associated data. H.G. and M.W.M.M conceived the project and wrote the manuscript.

## Data and materials availability

All sequencing data generated for this study have been deposited in NCBI’s Gene Expression Omnibus (Edgar, 2002) and are accessible through GEO Series accession number GSE169642 (PolII ChIP-sequencing and RNA-sequencing).

Published research reagents from the FMI are shared with the academic community under a Material Transfer Agreement (MTA) having terms and conditions corresponding to those of the UBMTA (Uniform Biological Material Transfer Agreement).

## Supplementary figures

**Figure S1:**
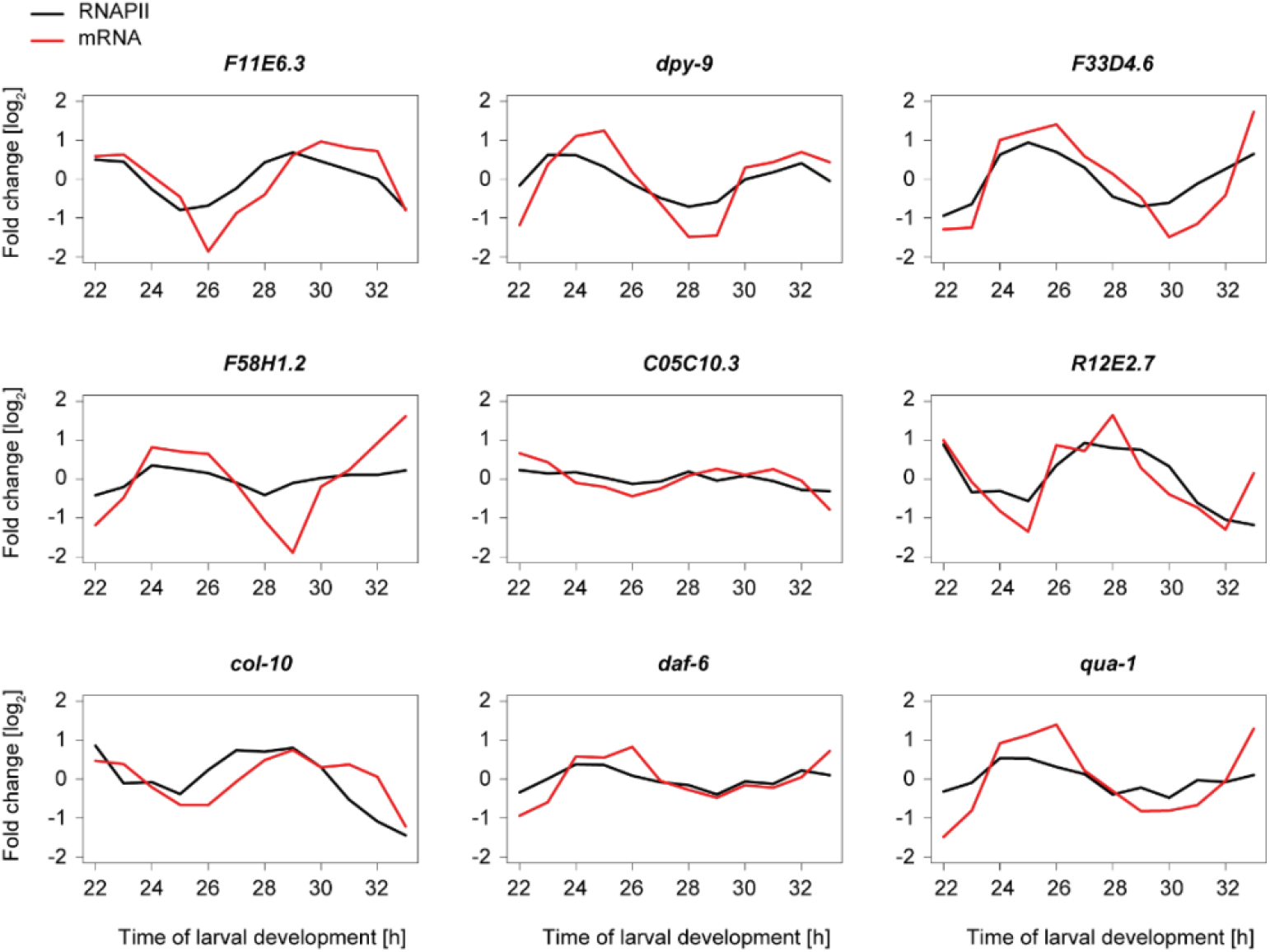
Comparison of RNAP II binding and mRNA level dynamics for specific genes, Related to Figure 1. Comparison of RNAPII ChIPseq and RNA-seq patterns over time, for the same samples, for genes for which we also established promoter-based reporter transgenes. With the exception of *C05C10*.*3* and *F58H1*.*2*, we can observe oscillatory oscillations, mostly preceding the mRNA transcript oscillation.

**Figure S2:**
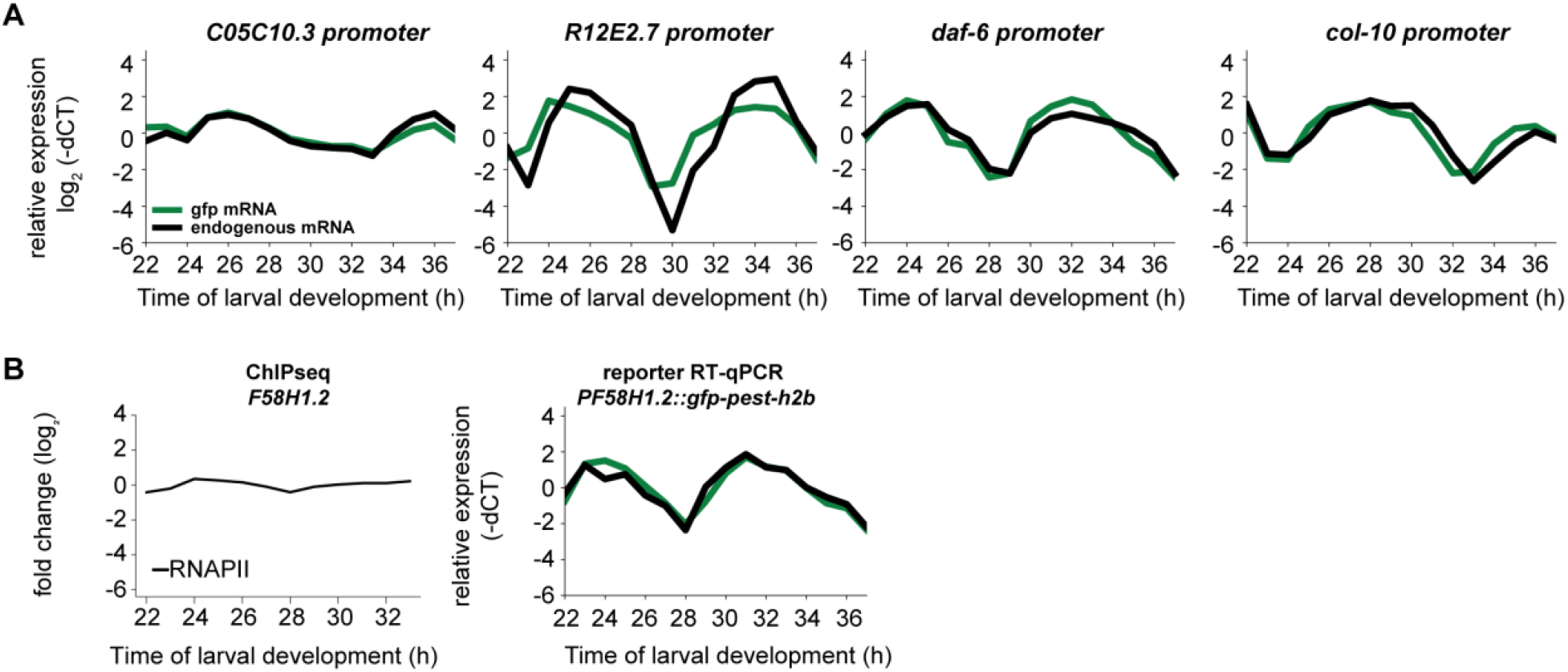
Additional transcriptional reporters investigated by RT-qPCR time courses; Related to Figure 1. **A**, Four additional transcriptional reporters were assayed by RT-qPCR time courses. All except the reporter for *R12E2*.*7* recapitulated the amplitude and the peak phase. In the *R12E2*.*7* case, we assume that we did not capture the entire promoter sequence or the effect of a distant regulatory element. **B**, Comparison of ChIP-seq reads (left) and RT-qPCR reporter gfp and endogenous levels of *F58H1*.*2*. We detect a large amplitude in the RT-qPCR experiment even though the amplitude is low in the ChIP-seq experiment.

**Figure S3:**
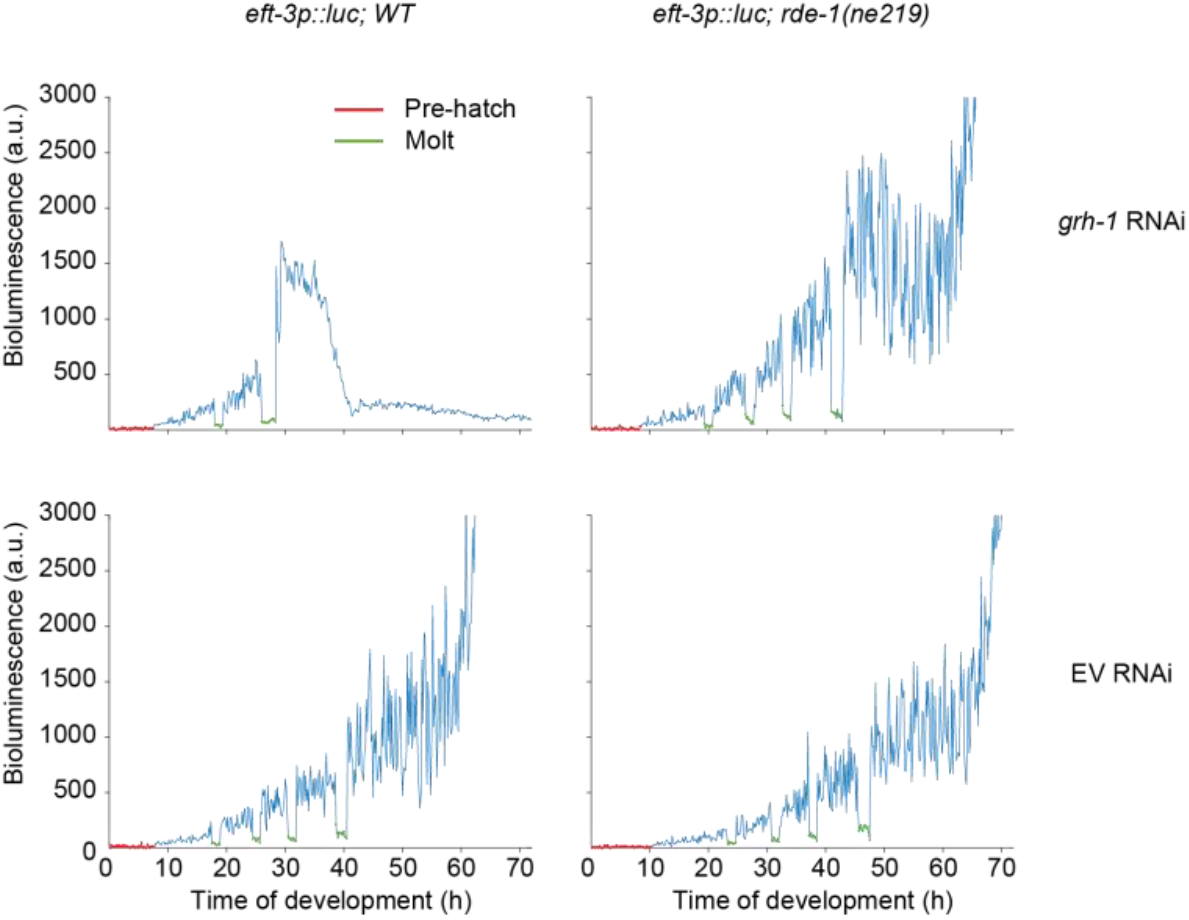
*grh-1* RNAi causes aberrant progression through development, Related to Figure 2. Representative raw luminescence trace of single animals grown at 20°C for 72 hours. Left column showing wildtype animals and right column RNAi-deficient animals (*rde-1(ne219)*). Animals in upper panel were grown in the presence of *grh-1* RNAi and animals in the lower panel were grown in the presence of mock (empty vector) RNAi. Pre-hatch is indicated in red and molts in green.

**Figure S4:**
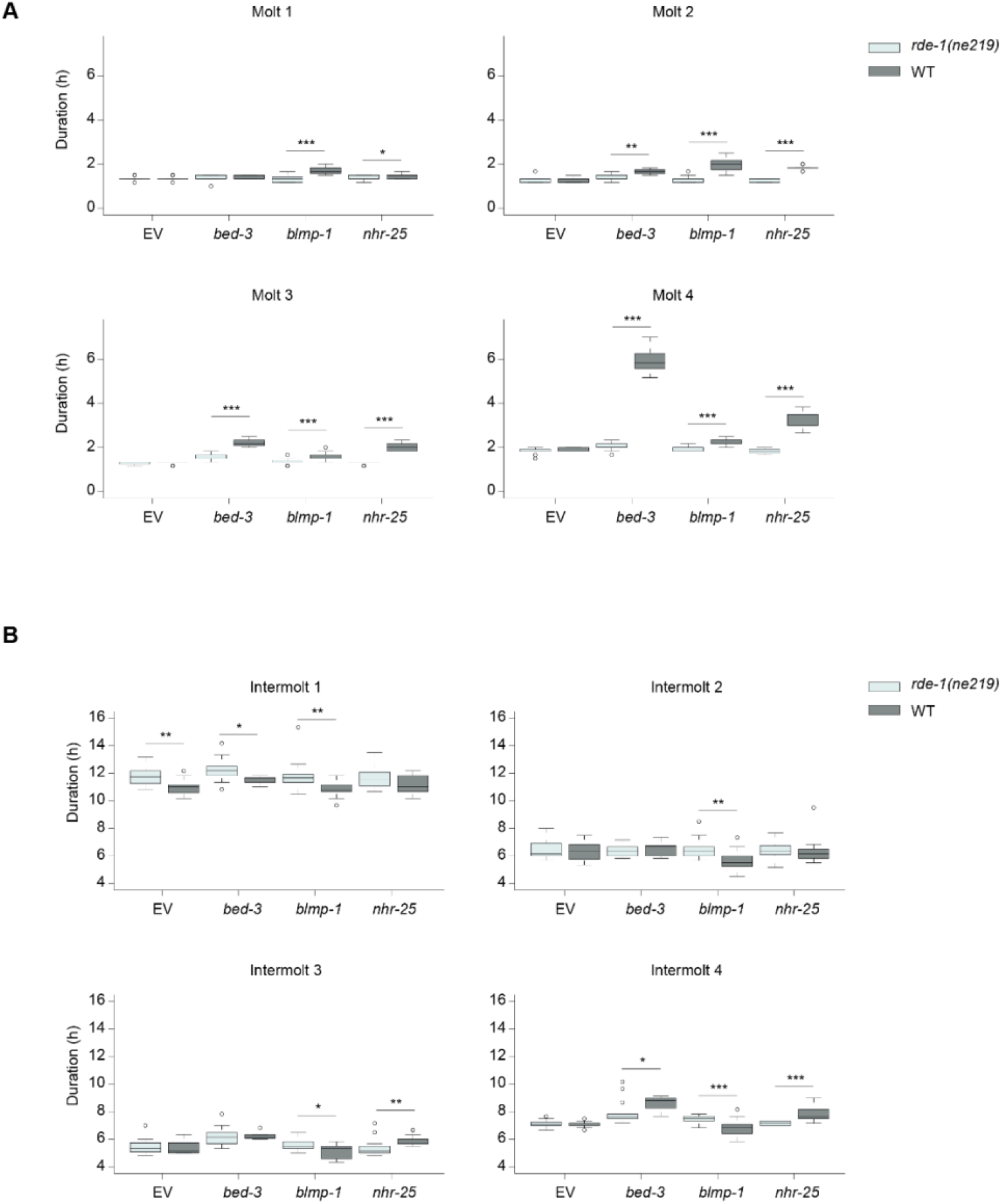
Quantification of molt and intermolt duration in screen hits that complete development, Related to Figure 2. **A**, Quantification of molt durations in RNAi deficient (*rde-1(ne219)*) animals (light grey) grown in the presence of mock (n=16), *bed-3* (n=14), *blmp-1* (n=16) or *nhr-25* (n=15) RNAi, and wildtype animals (dark grey) grown in the presence of mock (n=16), *bed-3* (n=8), *blmp-1* (n=16) or *nhr-25* (n=13) RNAi. Significant differences relative to *rde-1(ne219)* for each RNAi condition are indicated. **B**, Quantification of intermolt durations in RNAi deficient (*rde-1(ne219)*) animals (light grey) grown in the presence of mock (n=16), *bed-3* (n=14), *blmp-1* (n=16) or *nhr-25* (n=15) RNAi, and wildtype animals (dark grey) grown in the presence of mock (n=16), *bed-3* (n=8), *blmp-1* (n=16) or *nhr-25* (n=13) RNAi. Significant differences relative to *rde-1(ne219)* for each RNAi condition are indicated. * p<0.05, ** p<0.01, *** p<0.001, Wilcoxon test.

**Figure S5:**
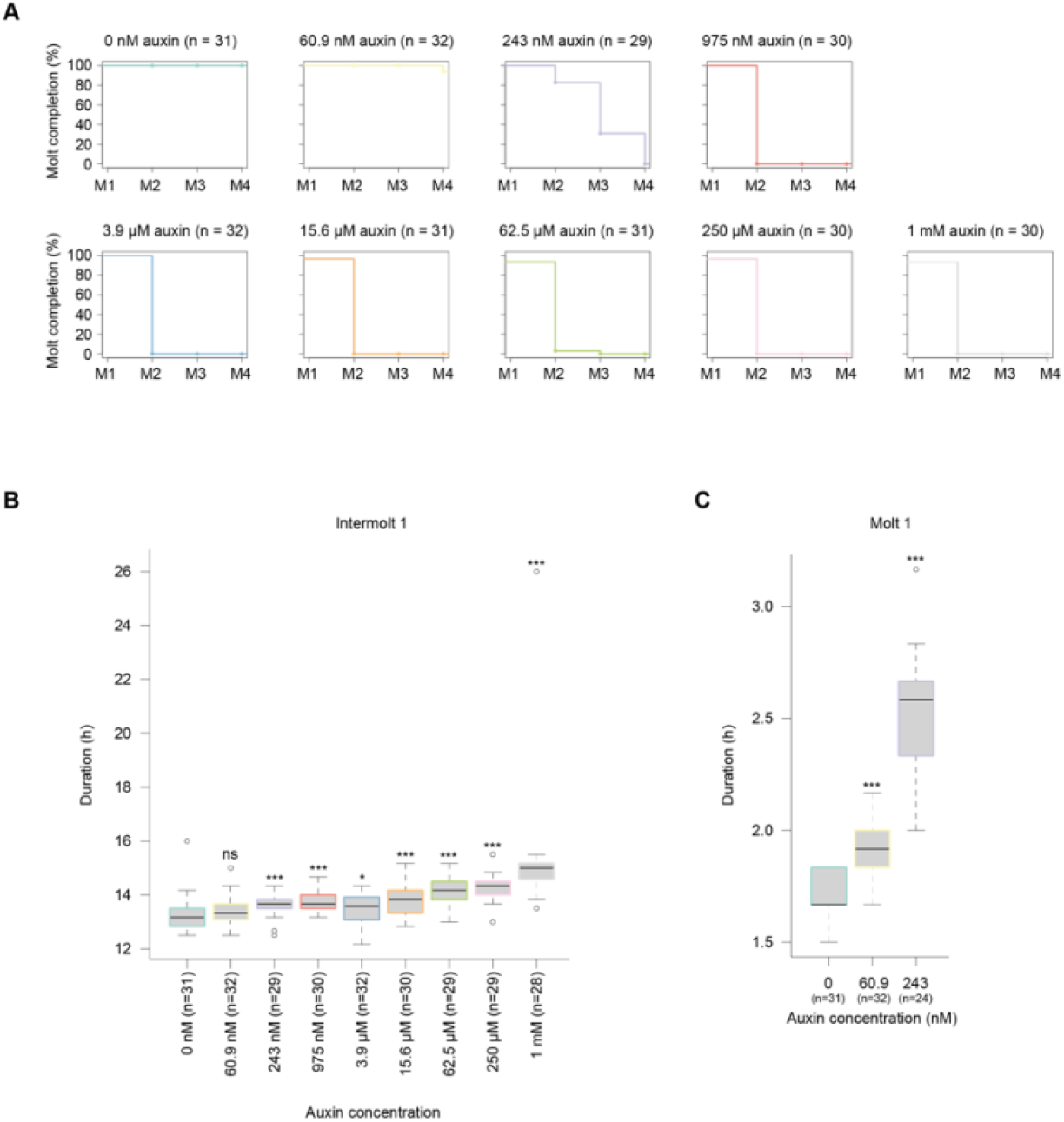
Intermolt is lengthened upon GRH-1 depletion at higher auxin concentrations, Related to Figure 4. **A**, Quantification of the percentage of *grh-1(xe135); eft-3p::luc; eft-3p::TIR1* animals (HW2434) that enter each of four molts molt upon increasing concentrations of auxin. Note that for some concentrations M1 does not start at 100%, i.e. animals initiated I1 but M1 was not observed. **B**, Boxplot showing the duration of the first intermolt of animals treated with indicated concentrations of auxin. Animals that failed to progress development beyond M1 (**A**). Significant differences relative to 0 nM auxin are indicated. **C**, Boxplot showing the duration of M1 of animals treated with indicated concentrations of auxin. Animals that failed to progress development beyond M1 were excluded (**A**). Significant differences relative to 0 nM auxin are indicated. P-values were determined by Wilcoxon test. ns: not significant, * p<0.05, ** p<0.01, *** p<0.001

**Figure S6:**
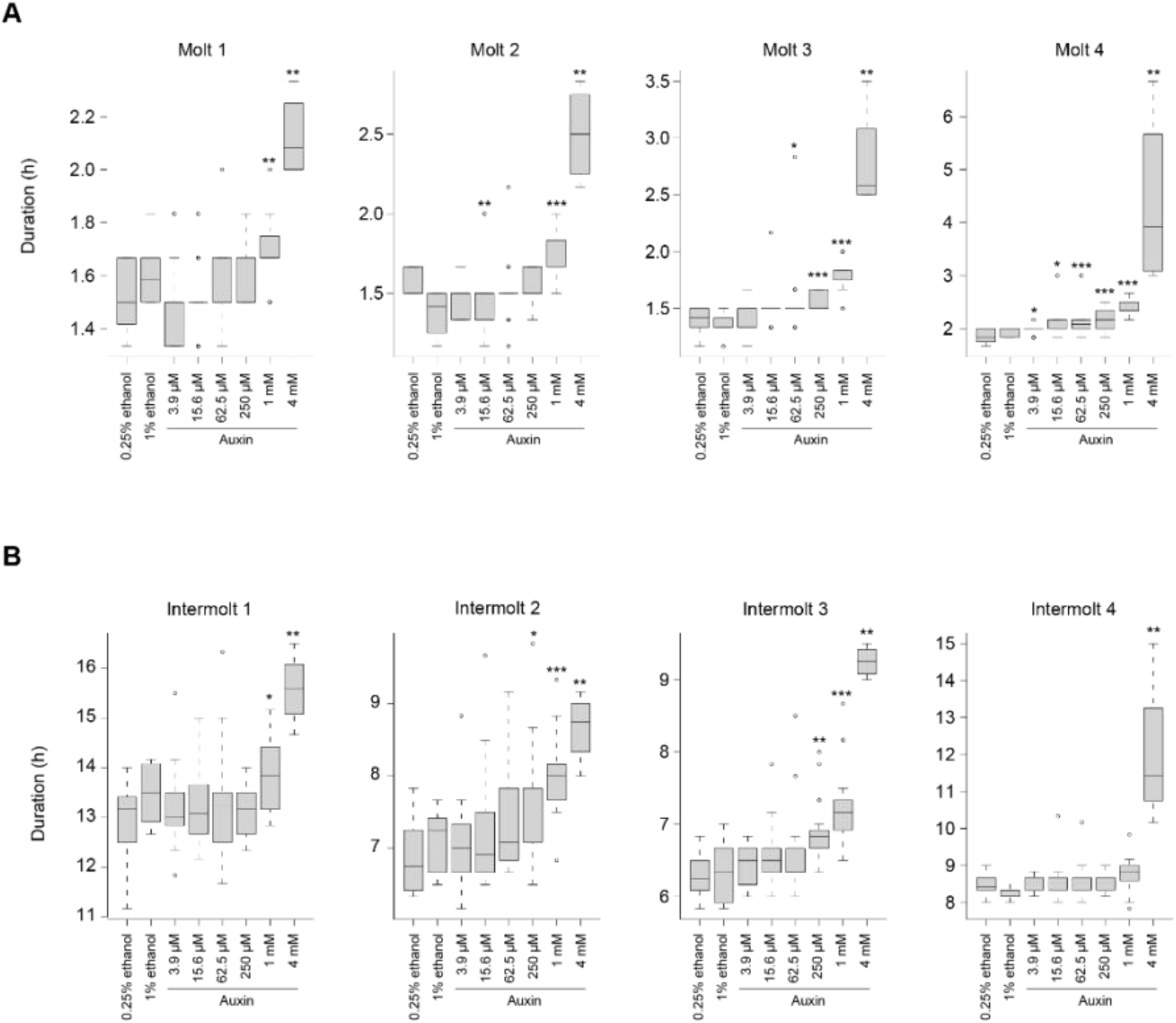
Effects of auxin in animal lacking a degron, Related to Figure 4. A, Boxplots showing duration of each molt for indicated concentrations of auxin in *eft-3p::TIR1; eft-3p::luc* animals (HW1984), which lack the *aid* tag on *grh-1*. Differences relative to 0.25% ethanol (0 nM auxin) for 3.9 μM – 1mM auxin and relative to 1% ethanol (0 nM auxin) for 4 mM auxin are indicated (ns: not significant, * p<0.05, ** p<0.01, *** P<0.001, Wilcoxon test). Note that concentrations ≤ 250 μM yield at most minor extensions of molt durations (compare Figure 3). At 4 mM auxin, most animals fail to complete M4. B, Boxplots showing duration of each intermolt for indicated concentrations of auxin in *eft-3p::TIR1; eft-3p::luc* animals (HW1984). Differences relative to 0.25% ethanol (0 nM auxin) for 3.9 μM-1mM auxin and relative to 1% ethanol (0 nM auxin) for 4 mM auxin are indicated (ns: not significant, * p<0.05, ** p<0.01, *** P<0.001, Wilcoxon test). Note that concentrations ≤ 250 μM yield at most minor extensions of intermolt durations.

**Figure S7:**
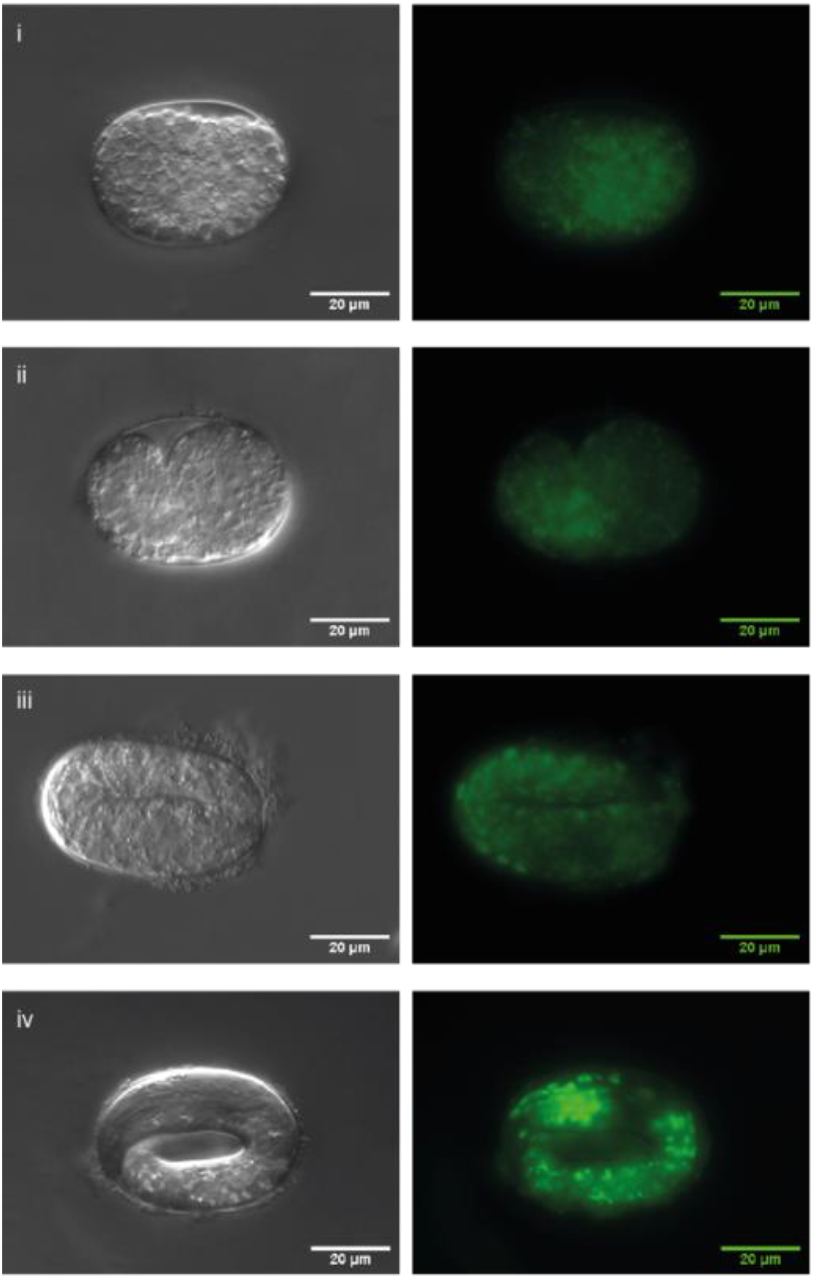
GFP-GRH-1 becomes detectable in elongating embryos, Related to Figure 6. Micrographs of *gfp::grh-1* embryos, order by increasing age top to bottom, captured by DIC (left) and fluorescence microscopy (right).

**Figure S8:**
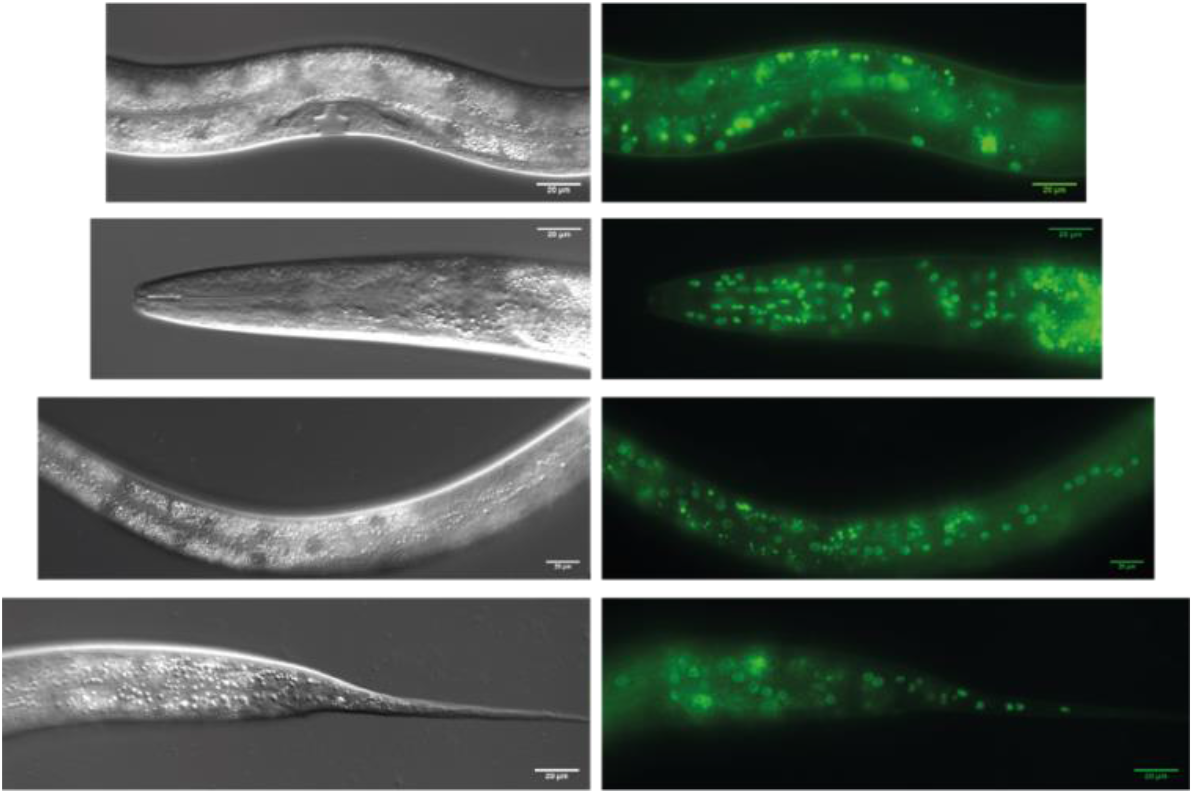
GFP-GRH-1 accumulates in various tissues during larval development, Related to Figure 6. DIC (left) and fluorescence microscopy (right) micrographs of *gfp::grh-1* larvae reveals *grh-1* expression in various stages

**Figure S9:**
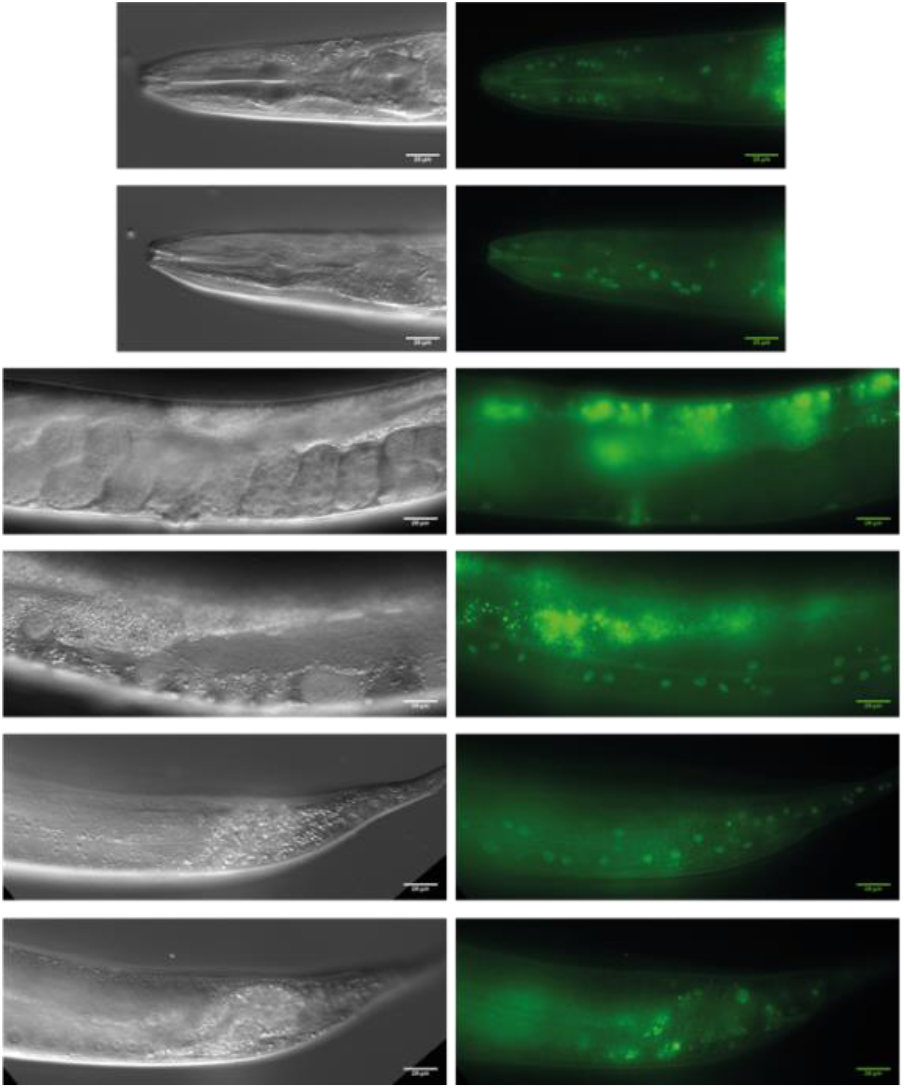
GFP-GRH-1 accumulation is greatly reduced in adults, Related to Figure 6. DIC (left) and fluorescence microscopy (right) micrographs of *gfp::grh-1* gravid adults reveals reduced *grh-1* expression levels in various tissues.

**Figure S10:**
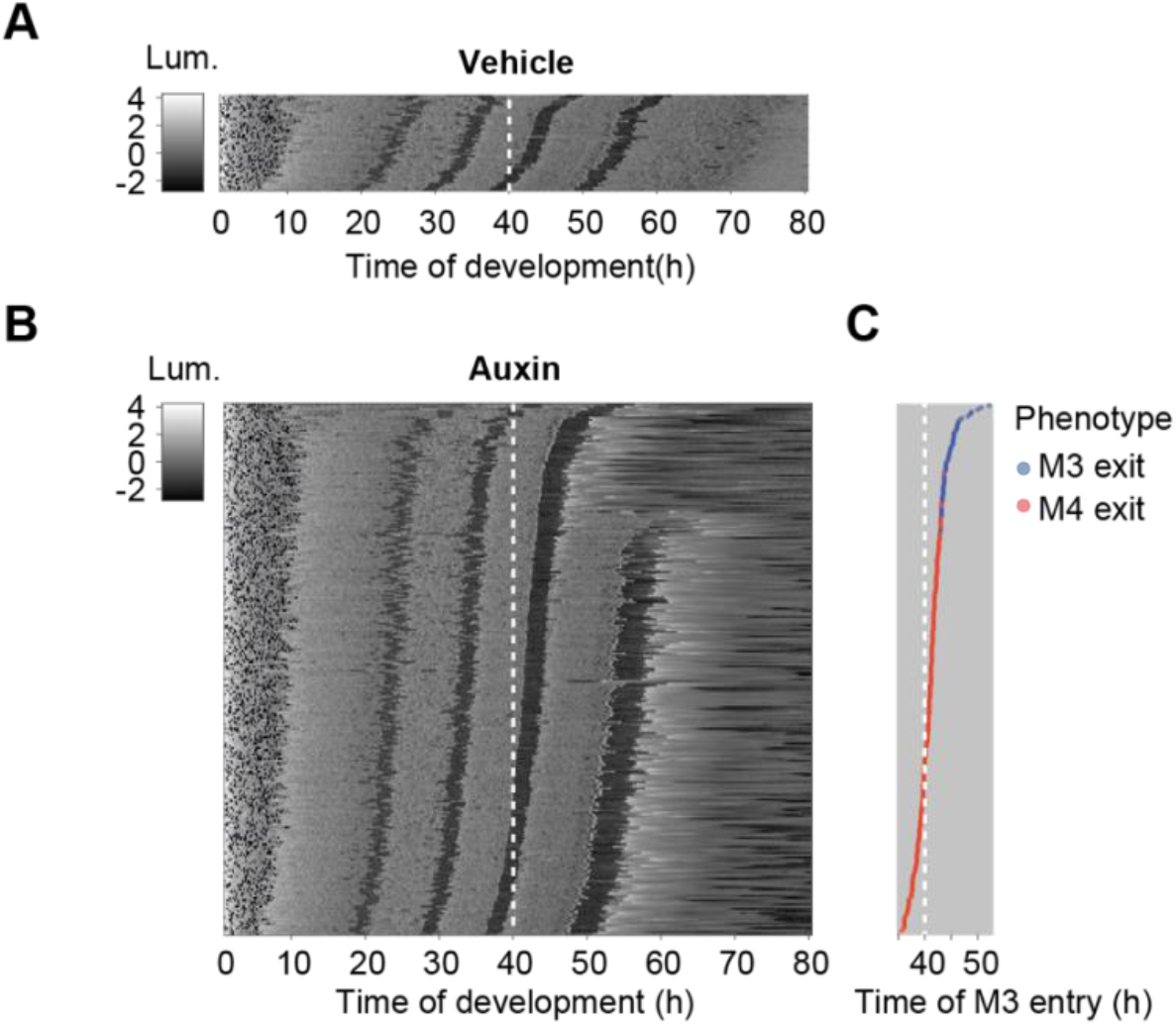
Molt exit phenotype is dependent on the time of GRH-1 depletion, Related to Figure 7. **A**,**B**, Heatmap showing trend-correct luminescence (Lum) of *grh-1(xe135); eft-3p::luc; eft-3p::TIR1* animals (HW2434) treated with vehicle (0.25% ethanol, A) or 250 μM auxin (B) at 40 hours (white dashed line). Black intensities correspond to low luminescence occurring during lethargus (molt). Embryos of various stages were left to hatch and develop during the assay. Luminescence traces are sorted by entry into molt 3 (M3) such that traces of early hatched animals are at the bottom and those of late hatched animals are at the top. **C**, Quantification of time at which animals enter molt 2 (M2 entry). Animals were assigned to M3 exit phenotype (blue) or M4 exit phenotype (red) when M3 or M4 was the last observed molt in the luminescence trace respectively. Animals sorted according to B.

